# A zero-agnostic model for copy number evolution in cancer

**DOI:** 10.1101/2023.04.10.536302

**Authors:** Henri Schmidt, Palash Sashittal, Benjamin J. Raphael

## Abstract

**Motivation:** New low-coverage single-cell DNA sequencing technologies enable the measurement of copy number profiles from thousands of individual cells within tumors. From this data, one can infer the evolutionary history of the tumor by modeling transformations of the genome via copy number aberrations. A widely used model to infer such *copy number phylogenies* is the *copy number transformation* (CNT) model in which a genome is represented by an integer vector and a copy number aberration is an event that either increases or decreases the number of copies of a contiguous segment of the genome. The CNT distance between a pair of copy number profiles is the minimum number of events required to transform one profile to another. While this distance can be computed efficiently, no efficient algorithm has been developed to find the most parsimonious phylogeny under the CNT model.

**Results:** We introduce the *zero-agnostic copy number transformation* (ZCNT) model, a simplification of the CNT model that allows the amplification or deletion of regions with zero copies. We derive a closed form expression for the ZCNT distance between two copy number profiles and show that, unlike the CNT distance, the ZCNT distance forms a metric. We leverage the closed-form expression for the ZCNT distance and an alternative characterization of copy number profiles to derive polynomial time algorithms for two natural relaxations of the small parsimony problem on copy number profiles. While the alteration of zero copy number regions allowed under the ZCNT model is not biologically realistic, we show on both simulated and real datasets that the ZCNT distance is a close approximation to the CNT distance. Extending our polynomial time algorithm for the ZCNT small parsimony problem, we develop an algorithm, *Lazac*, for solving the large parsimony problem on copy number profiles. We demonstrate that *Lazac* outperforms existing methods for inferring copy number phylogenies on both simulated and real data.

**Availability:** *Lazac* is implemented in C++17 and is freely available at: github.com/raphael-group/lazac-copy-number.

## 1 Introduction

Tumor evolution is characterized by both small and large genomic alterations that alter the fitness of cancer cells [31]. *Copy number aberrations*, i.e. modifications to the number of copies of a genomic segment, are an important and frequent sub-class of such alterations that drive prognostic and metastatic outcomes [1]. Deriving the evolutionary history of copy number aberrations, herein referred to as *copy number phylogenies*, is thus important for understanding the emergence of primary tumors and the development of subpopulations of cells that evade treatment and/or metastasize to other anatomical sites.

Recent technological and computational improvements in single-cell sequencing have enabled the mapping of high resolution copy number profiles in single cells. For example, the high-throughput 10x Genomics Single-cell Copy Number Variation solution [2, 48] produces ultra-low coverage (< 0.05×) whole genome sequencing data from ≈ 2000 individual cells. Other recent technologies, including DLP/DLP+ [49, 23, 15] and ACT [28], produce similar data. Multiple computational methods [48, 45, 44, 10, 20, 24] have been introduced to infer high resolution *copy number profiles*, integer vectors that contain the number of copies of each genomic segment, from this type of data. Other recent methods can infer copy number profiles from thousands of cells or spatial locations from single-cell RNA sequencing (scRNA-seq) [16], scATAC-seq [47], or spatial transcriptomics data [12].

The increasing availability of technologies to measure genomic copy number in thousands of cell motivates development of methods to infer the cellular phylogenies from copy number profiles. However, there are multiple challenges in inferring phylogenies from copy number profiles. First, copy number aberrations are diverse, ranging from small duplications and deletions [3] to whole chromosome shattering and reconstruction events [41]. Second, a single copy number aberration can alter the number of copies of a large section of the genome *simultaneously*. This means that loci on the genome cannot be treated as independent phylogenetic characters, a widely-used assumption in phylogenetics [18, 19, 39, 46] Finally, the increasing size (> 10, 000 cells) and resolution (< 5Kb bins) of copy number profiles require increasingly scalable algorithms.

One widely used model of copy number evolution is the *copy number transformation* (CNT) model [38]. In the CNT model, a genome is represented as a vector of non-negative integers and copy number aberrations correspond to the increase or decrease of the entries in a contiguous *interval* of coordinates in the vector, explicitly modeling the non-independence of copy number amplifications and deletions. The *CNT distance* is the minimum number of copy number events needed to transformation one profile to another. The CNT distance is computable in linear time [51] and has been used to define an evolutionary distance between profiles. Since the CNT distance is not symmetric, a variety of symmetrized CNT distances have also been used to construct copy number phylogenies using distance-based phylogenetic methods [38, 11, 50, 22]. Further, owing to its effectiveness, the CNT model has become the basis of a variety of distinct models [50, 22, 8] for copy number evolution.

While the CNT model is described by specific events – or mutations – there has been little work on constructing phylogenetic trees under the CNT model using the method of maximum parsimony. Even the small parsimony problem – where the topology of the tree is given and one aims to infer the ancestral profiles that minimizes the total number of copy number events on the tree – has no known efficient solution. For example, for the special case of a two leaf tree, the best algorithm for the CNT small parsimony problem [51] runs in 𝒪(*nB*^7^) time where *B* is the largest allowed copy number and *n* is the number of loci [11]. Without an efficient algorithm for the small parsimony problem under the CNT model, one cannot hope to solve the large parsimony problem, where the topology of the tree is unknown.

We introduce a relaxation of the CNT model, called the *zero-agnostic copy number transformation* (ZCNT) model, that approximates the CNT model and has a number of desirable properties. Unlike the CNT model, the ZCNT model allows for the amplification of zero copy number regions. While such an operation is not biologically realistic, we show that this relaxation makes the ZCNT distance a metric, in contrast to the CNT distance. Moreover, we derive a closed form expression for the ZCNT distance between two profiles. We use this closed form expression as well as an alternative characterization of copy number profiles to solve two relaxations of the small parsimony problem in polynomial time. To our knowledge, this is the first attempt to solve the small parsimony problem for a segment-based (i.e. non-independent) model of copy number evolution. We then use our efficient algorithm for the (relaxed) small parsimony problem to design an algorithm, *Lazac* (Large-scale Analysis of Zero Agnostic Copy number), for inferring copy number phylogenies by solving the large parsimony problem. We show on simulated data that *Lazac* is > 100× faster than other phylogenetic methods and also more accurate in recovering the ground truth phylogeny. On single-cell whole-genome sequencing data from human breast and ovarian tumors, *Lazac* finds phylogenies that are more consistent with both copy number clones and single-nucleotide variants (SNVs).

## 2 Copy number transformations

A *copy number profile p* = [*p*_1_, …, *p*_*m*_] is a vector of non-negative integers where *p*_*j*_ ∈ {0, …, *B*} is the the number of copies of locus *j*. Suppose we measure the copy number profile of *n* cells of a tumor across *m* loci in a single-cell DNA sequencing experiment. We encode the copy number profiles in a *n* × *m copy number matrix M* = [*M*_*i,j*_] where *M*_*i,j*_ ∈ {0, …, *B*} is the copy number of cell *i* at locus *j*. The copy number profile *p* of cell *i* is then the *i*^th^ row of this matrix, and is denoted *M*_*i*_.

One of the most basic phylogenetic principles is that nearly perfect measurement of evolutionary distances enables exact recovery of the evolutionary history [40]. It is thus not surprising that many of the successful attempts at inferring copy number phylogenies focus on finding *good* methods to compute an evolutionary distance between a pair of copy number profiles. Early methods for computing evolutionary distances on copy number data [3, 7] employed simple measures of distance such as the Hamming, weighted Hamming, and *ℓ*_1_ distance between copy number profiles. However, these distances do not account for dependencies between loci caused by long CNAs spanning contiguous segments of the genome, leading to inaccurate phylogenetic reconstruction [38, 22].

In this section, we describe and investigate the copy number transformation (CNT) model, one of the most well-known and successful evolutionary models for copy number evolution in cancer. The CNT model was originally introduced in MEDICC [38] and extended in subsequent studies [51, 11, 50, 8, 22]. Since the CNT model only allows intrachromosomal copy number events, it is sufficient to consider the case of a single chromosome, and thus for ease of exposition we will describe the model using a single chromosome.

The fundamental operation in the CNT model is a *copy number event* which increases or decreases (by one) the entries in a contiguous interval of a copy number profile, defined formally as follows.

### Definition 1

(Copy number event). *A* copy number event 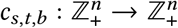 *is a function that maps a copy number profile* 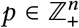 *to a profile c*_*s,t,b*_ (*p*) *described by its entries as*

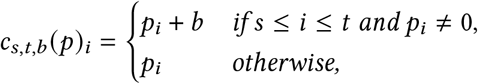

*where s* ≤ *t and b* ∈ {+1, −1}. *We denote such a function as c when clear by context*.

That is, an amplification (resp. deletion) increases (resp. decreases) the copy number of all *non-zero* entries in the interval between positions *s* and *t*, or alternatively a copy number event *skips* the zero entries (Figure 1). Thus, once a locus is lost (i.e. *p*_*i*_ = 0), the locus cannot be regained or deleted further. A *copy number transformation C* is the composition of multiple copy number events and we denote this function as *C* = (*c*_1_, …, *c*_*n*_) when *C*(*p*) = *c*_*n*_ (⋯ (*c*_2_(*c*_1_(*p*)))).

**Figure 1:**
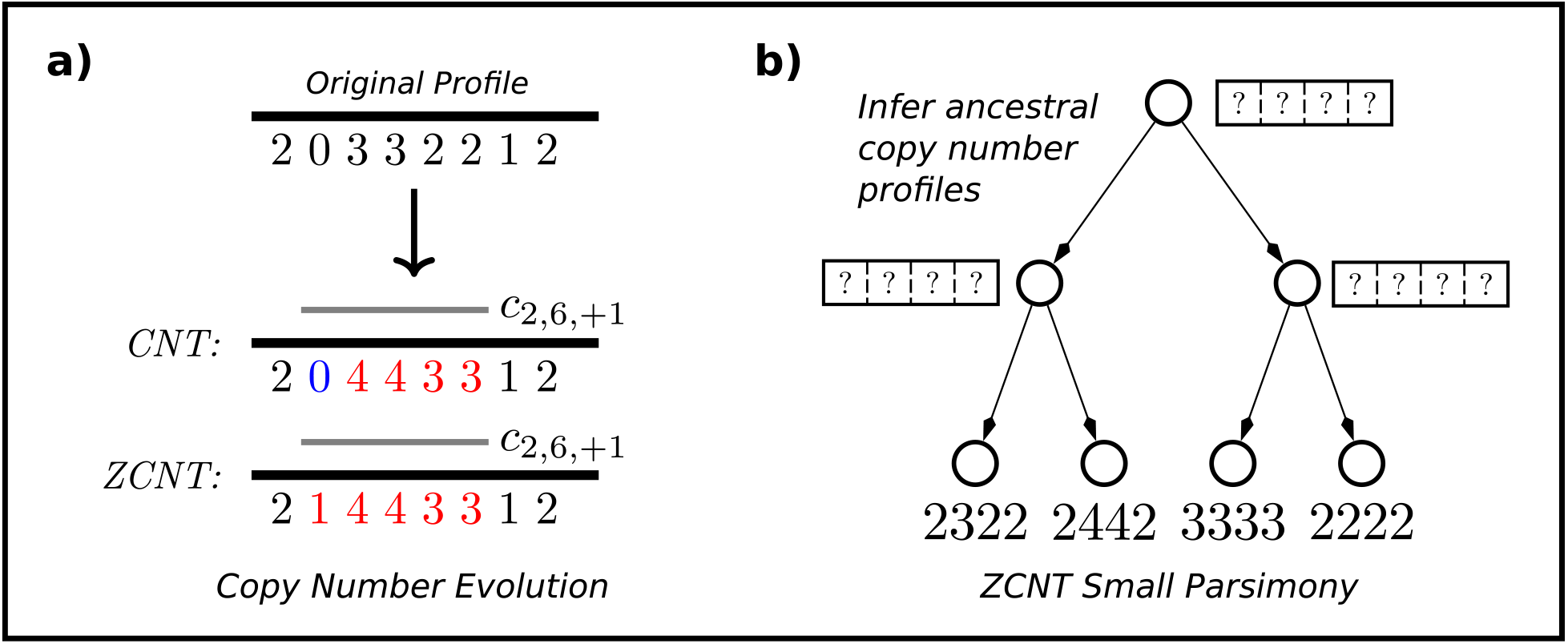
**(a)** The results of applying a copy number transformation *c*_2,6,+1_ under both the CNT model and ZCNT model. Under the CNT model a zero cannot be increased via the amplification, but the zero-agnostic CNT model allows the zero to increase to one copy. **(b)** An instance of the ZCNT small parsimony problem: given a tree with copy number profiles labeling the leaves, the goal is to infer the ancestral copy number profiles that minimize the total ZCNT distance across all edges.

Several copy number problems have been previously studied to compute evolutionary distances under the CNT model. The first, and simplest, is the *copy number transformation problem*, originally introduced in [38], which defines a distance, *σ*(*u, v*), between two copy number profiles. Put simply, the distance between two profiles is the length of the shortest copy number transformation needed to transform one profile to another.

### Definition 2

(Copy number transformation distance). *Given two copy number profiles u and v, the* copy number transformation distance *is*

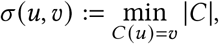

*where C* = (*c*_1_, …, *c*_*n*_) *is a CNT. Alternatively, σ*(*u, v*) = ∞ *if no such transformation exists*.

[51, 43] show there is a (non-trivial) strongly linear time algorithm (i.e. time complexity 𝒪(|*u*| + |*v*|)) for computing the CNT distance *σ*(*u, v*). Unfortunately, the CNT distance *σ*(*u, v*) is not symmetric (i.e. *σ*(*u, v*) ≠ *σ*(*v, u*)), which makes it difficult to use in distance based phylogenetic methods such as neighbor joining [34].

In order to apply distance-based phylogenetic methods, multiple approaches to symmetrize the distance *σ*(*u, v*) have been introduced. [51] use a mean correction replacing the asymmetric *σ*(*u, v*) with a symmetric distance *σ*′(*u, v*) defined as

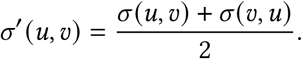

Alternatively, several authors [38, 22, 11] define the distance between two profiles in terms of a closely related, median profile *w*. Specifically, the *median distance* between two profiles *u* and *v* is defined to be the smallest value of

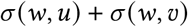

over all profiles *w*. Computing this median distance is called the *copy number triplet problem* in [11]. Unfortunately, no efficient algorithm is known for the copy number triplet problem. The fastest algorithm uses 𝒪(*mB*^7^) time and 𝒪(*mB*^4^) space where *B* is the maximum allowed copy number [11].

## 3 Small and large copy number parsimony

The small parsimony problem for copy number profiles is the following: given a tree 𝒯 whose leaves are labeled by copy number profiles, infer ancestral copy number profiles that minimize the total dissimilarity between profiles across all edges (Figure 1). For evolutionary models in which each character evolves independently and has finitely many states (e.g. single nucleotide substitution models), the small parsimony problem is solved in polynomial time via Sankoff’s algorithm, a dynamic programming algorithm [36]. Unfortunately, the CNT model presents two major challenges in solving the small parsimony problem. First, since copy number events affect multiple loci simultaneously, the loci cannot be analyzed independently, in contrast to most phylogenetic characters. Second, the space of possible copy number profiles is a priori unbounded, since the maximum copy number of a segment in a genome is unknown. Thus, it is not surprising that there is no published solution to the small parsimony problem for CNT dissimilarity, with the exception of the special case of two-leaf trees [11]. Here, we formalize both the CNT small parsimony problem and the corresponding large parsimony problem, the latter of which was previously described in [11].

A *copy number phylogeny*(𝒯, *ℓ*) is a rooted tree 𝒯 and leaf labeling *ℓ*. Let *V*(𝒯), *E*(𝒯), and *L*(𝒯) denote the edges, vertices, and leaves of 𝒯, respectively. In our applications below, each leaf of 𝒯 represents one of the *n* cells (or bulk samples) from a tumor. An ancestral labeling 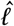 of a copy number phylogeny is a vertex labeling of 𝒯 that agrees with *ℓ* on the leaves of 𝒯, i.e. 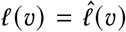 when *v* ∈ *L*(𝒯). We say that (𝒯, *ℓ*) is a copy number phylogeny for copy number matrix *M* if 𝒯 has *n* leaves such that *ℓ* labels each leaf by a row of *M*. Formally, if (𝒯, *ℓ*) is a copy number phylogeny for a copy number matrix *M*, then there exists a cell assignment *π* : [*n*] → *L*(𝒯) that assigns each cell *i* to a leaf *v* such that *ℓ*(*v*) = *M*_*i*_.

We define the cost 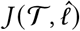 of a vertex labeled, copy number phylogeny as the total number of copy number events required to explain the phylogeny:

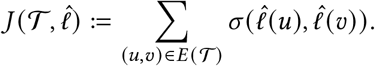

We now introduce the small parsimony problem [14] under the copy number transformation model.

### Problem 1

(CNT Small parsimony). *Given a copy number phylogeny* (𝒯, *ℓ*) *find an ancestral labeling* 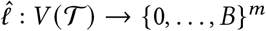 *such that (i)* 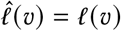 *for all leaves v* ∈ *L*(𝒯) *and (ii)* 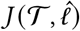 *is minimized*.

The *parsimony score* is defined as the cost 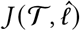 of the solution 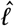 to the CNT small parsimony problem. To the best of our knowledge the CNT small parsimony problem (Problem 1) has not been analyzed in the literature. We believe this is due to the difficulty of solving the CNT small parsimony problem. That is, for even a special case of two-leaf trees, referred to as the *copy number triplet* problem [11], no strongly polynomial time algorithm is known (Section 2).

The CNT large parsimony problem defined in [11], aims to find a vertex labeled, copy number phylogeny 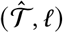 for a matrix *M* with minimum cost.

### Problem 2

(CNT Large parsimony). *Given copy number matrix M, find a copy number phylogeny* (𝒯, *ℓ*) *for M and ancestral labeling* 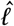 *such that* 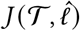 *is minimized*.

Unsurprisingly, [11] showed that the above large parsimony problem (Problem 2) is NP-hard. They also formulated an integer linear program (ILP) to solve the problem exactly. However, this ILP consists of *O*(*n*^2^*m* + *nm* log *B*) variables and does not scale to the size of current real data sets with thousands of cells.

## 4 The zero-agnostic CNT model

The copy number transformation (CNT) model imposes the constraint that once a locus is lost (has zero copy number), the locus remains with zero copies for all time. While this constraint is biologically realistic, the constraint also makes the inference problems – including the CNT small (and large) parsimony problems – computationally hard to solve. Here, we show that *relaxing* the constraint that copy number events do not alter zero entries leads to a simpler model with favourable mathematical properties. We call this the *zero-agnostic* copy number model (Figure 1) to indicate that the model allows the amplification and deletion of loci with zero copies. Formally, we define a *zero-agnostic copy number event* as follows.

### Definition 3

(Zero-agnostic copy number event). *A* zero-agnostic copy number event *c*_*s,t,b*_ : ℤ^*n*^ → ℤ^*n*^ *is a function that maps a profile p* ∈ ℤ^*n*^ *to a profile written c*_*s,t,b*_(*p*) *described by its entries as*

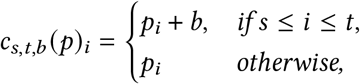

*where s* ≤ *t and b* ∈ {+1, −1}. *We denote such a function as c when clear by context*.

Thus, a zero-agnostic copy number event either increases or decreases the number of copies of all loci in the interval (*s, t*) regardless of whether the loci have zero copies. While our formulation allows for the number of copies of a locus to decrease below zero, one can show that given two profiles with non-negative entries, it is always possible to find an optimal ZCNT such that no intermediate profile has negative entries. Specifically, as a corollary to the commutativity of zero-agnostic copy number transformations (Proposition 2), one can re-order events such that amplifications always occur first.

Due to space constraints, we do not include all proofs in the main text. Any proof not present in the main text can be found in Supplementary Proofs C.

### 4.1 Delta profiles

We simplify our analysis of zero-agnostic copy number events by examining their effect on the differences between the copy number of adjacent loci. In particular, while a zero-agnostic copy number event *c*_*s,t,b*_ increments (or decrements) all entries *p*_*i*_ where *i* ∈ {*s*, …, *t*}, *c*_*s,t,b*_ only alters two *differences* between adjacent loci, namely the difference *p*_*s*_− *p*_*s* −1_ and the difference *p*_*t*+1_ − *p*_*t*_. To formalize this idea, we first define the *delta profile*, vectors obtained by taking the differences in copy number between adjacent loci.

#### Definition 4

*A* delta profile *is any vector p* ∈ ℤ^*n*^ *that satisfies the* balancing condition:

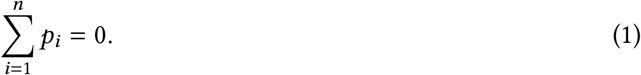

*Or equivalently*, 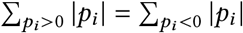. *We denote the set of delta profiles in ℤ*^*n*^ *as* 𝒟_*n*_.

The above definition provides us with a convenient (and useful) description of the image of the following difference transformation, which we call the *delta map*.

#### Definition 5

*The delta map* Δ : ℤ^*n*^ → 𝒟_*n*+1_ *maps a copy number profile p to a delta profile* Δ(*p*) *by taking the differences in adjacent copy number loci. Specifically*,

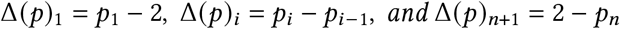

*where the constant* 2 *represents a normal, diploid copy number*.

A basic property of the delta map Δ : ℤ^*n*^ → 𝒟_*n*+1_ is that it is invertible.

#### Proposition 1

*The delta map* Δ : ℤ^*n*^ → 𝒟_*n*+1_ *is invertible*.

Since Δ is one-to-one and onto with respect to 𝒟_*n*+1_, each delta profile *p*′ then corresponds to a unique copy number profile *p* = Δ^−1^(*p*′).

Interestingly, a copy number event *c*_*s,t,b*_ applied to a copy number profile *p* only affects two entries of the delta profile Δ(*p*), meaning that loci of the corresponding delta profile are (nearly) independent. We formalize this in the following definition of a *delta event*.

#### Definition 6

(Delta event). *A* delta event *δ*_*s,t,b*_ : 𝒟_*n*_ → 𝒟_*n*_ *is a function that maps a delta profile p* ∈ 𝒟_*n*_ *to a delta profile δ*_*s,t,b*_ (*p*) *described by its entries as*

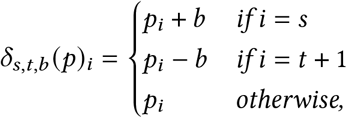

*where s* ≤ *t and b* ∈ {+1, −1}. *We denote such a function as δ when clear by context*.

A *delta transformation D* = (*δ*_1_, …, *δ*_*n*_) is the composition of multiple delta events, where *D*(*p*) = *δ*_*n*_(⋯ (*δ*_2_ (*δ*_1_ (*p*)))). We now state the connection between delta events and zero-agnostic copy number (ZCNT) events in the following theorem and corollary.

#### Theorem 1

*Let c*_*s,t,b*_ *be a zero-agnostic copy number event and δ*_*s,t,b*_ *be a delta event. Then*,

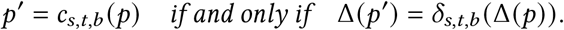

#### Corollary 1

*Let* 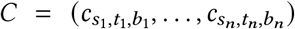 *be a zero-agnostic copy number transformation and* 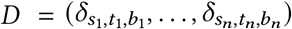 *be the corresponding delta transformation. Then*,

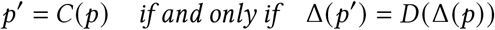

*Proof*. The corollary follows by induction on |*C*| and repeated application of (Theorem 1).

### 4.2 Computing the ZCNT distance

Let *d*(*u, v*) be the minimum number of *zero-agnostic* copy number events needed to transform the copy number profile *u* to *v*. In this section we derive a closed form expression for *d*(*u, v*).

We begin by noting that *d*(*u, v*) is equal to the minimum number *d*′(Δ(*u*), Δ(*v*)) of delta events needed to transform delta profile Δ(*u*) to Δ(*v*). This follows from the equivalence between the copy number transformations and the corresponding delta transformation (Corollary 1). Thus, it suffices to only consider delta profiles and delta events; for the rest of the section all profiles *u* and *v* are delta profiles unless otherwise specified.

We start by observing two basic facts: delta transformations are commutative and *d*′(*u, v*) forms a metric.

#### Proposition 2

*A delta transformation D* = (*δ*_1_, …, *δ*_*n*_) *is commutative. That is, the application of D to a profile is identical to the application of* 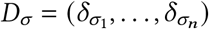 *where σ is any permutation of* {1, …, *n*}.

#### Proposition 3

*d*′(*u, v*) *is a distance metric. Further*,

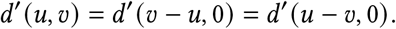

Note that this also implies that zero-agnostic copy number transformations are commutative and that *d*(·,·), is a distance metric. To see this, let 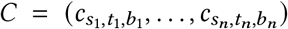 be a zero-agnostic copy number transformation and 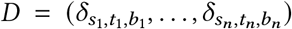 be the corresponding delta transformation, then for any vector *p* and permutation *σ*,

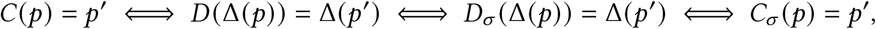

where the first equivalence follows from Corollary 1, the second from Proposition 2, and the third from Corollary 1. This implies that *C*(*p*) = *C*_*σ*_ (*p*), which proves that a zero-agnostic copy number transformation is commutative. To see that *d*(·, ·) is a distance metric, it suffices to observe that *d*(*u, v*) = *d*′(Δ(*u*), Δ(*v*)) implies symmetry and reflexivity. Then, the triangle inequality is satisfied since the composition of a zero-agnostic copy number transformation from *u* to *w* and *w* to *v* yields a copy number transformation from *u* to *v*.

From our characterization of delta profiles, we derive our expression for the distance between delta profiles.

#### Theorem 2

*For delta profiles u and v*, 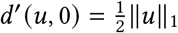. *Thus*, 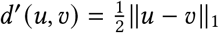.

*Proof*. Since each event decreases the total magnitude of ∥Δ(*u*)∥_1_ by at most two, to transform Δ(*u*) to the 0 profile requires at least 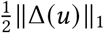 events.

We prove the other direction by induction on 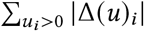. Clearly, if the sum is zero, the claim holds. Otherwise, by (Proposition 1), we can choose *c* to be any event that decrements *i* ∈ {*i* : Δ(*u*)_*i*_ > 0} and increments *j* ∈ {*j* : Δ(*u*)_*j*_ < 0}. Applying *c* to Δ(*u*) results in a delta profile Δ(*u*′) such that ∥Δ(*u*′)∥_1_ = ∥Δ(*u*)∥_1_ − 2. Invoking the induction hypothesis then yields a sequence of 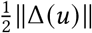 events to transform Δ(*u*) to the 0 profile.

The second statement follows from Proposition 3.

As a corollary to the above theorem and the equivalence between zero-agnostic copy number transformations and delta transformations (Corollary 1), we have our closed form expression for the ZCNT distance between copy number profiles.

#### Corollary 2

*For copy number profiles p and p*′,

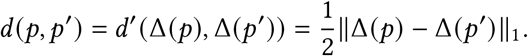

Further, as a corollary to the fact that *d*(*u, v*) is a distance metric, the following median distance is trivially computed in linear time:

#### Corollary 3

*Given two copy number profiles u and v, both u and v minimize the median distance d*(*w, u*) + *d*(*w, v*) *over all choices of copy number profiles w*. *Thus*,

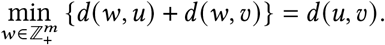

## 5 Algorithms

### 5.1 ZCNT small parsimony

We show below that the special form of the ZCNT model enables us to solve two natural relaxations of the small parsimony problem in polynomial time. First, using the equivalence between copy number profiles and delta profiles described above, we formulate the small parsimony problem (Problem 1) using the ZCNT model as follows.

#### Problem 3

(ZCNT Small Parsimony). *Given a copy number matrix M, a tree* 𝒯 *and cell assignment π* : [*n*] → *L*(𝒯), *find a vertex labeling ℓ* : *V*(𝒯) → ℤ^*m*^ *minimizing*

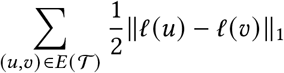

*such that the following two conditions are satisfied:*

i. *ℓ*(*π*(*i*)) = Δ(*M*_*i*_) *for all cells i* ∈ [*n*],
ii. *l*(*u*) *satisfies the balancing condition* (1) *for all vertices u* ∈ *V*(𝒯).

To solve the above problem, we recall the general form of the Sankoff-Rousseau recurrence [9, 36] for solving the small parsimony problem. Let *c*(𝒯; *x*) be the cost of the optimal labeling 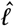 of *V*(𝒯) that agrees with copy number matrix *M* and has label *x* for the root. Let 𝒯_*w*_ denote the sub-tree rooted at *w* and suppose that *w* has children *u* and *v*. Then, by condition (ii), and the requirement that the ancestral labeling lies in ℤ^*m*^, we have the following recurrence relation [9]:

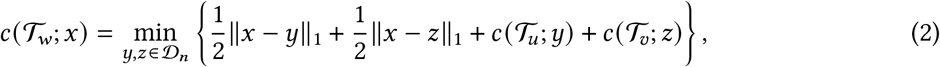

This recurrence has several difficulties. First, *y* and *z* are unbounded and can take on any value in ℤ^*m*^. Thus, it is impossible to store a dynamic programming table for *c*(*T*_*w*_; *x*) without imposing bounds on the maximum copy number. Further, even when the entries are constrained to a bounded interval {0, …, *B*}^*m*^, the dynamic pr ogrammin g table has size (*B* + 1)^*m*^, exponentially large. Second, because of the balancing conditions (1), 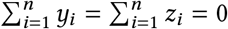, one cannot analyze the loci independently.

Despite these challenges, the recurrence (2) is a substantial improvement over the analogous recurrence under the CNT model. In fact, if we remove *either the balancing* (1) *or the integrality condition*, we can solve this recurrence in (resp. strong or weak) polynomial time.

#### Theorem 3

*If the balancing condition* (1) *is dropped, the ZCNT small parsimony problem can be solved in* 𝒪(*mn*) *time. If the integrality condition is dropped, the ZCNT small parsimony problem can be solved in (weakly) polynomial time using a linear program with* 𝒪(*mn*) *variables and constraints*.

Both of these facts derive from our closed form expression for the ZCNT distance between two copy number profiles in terms of the *ℓ*_1_ norm. We sketch the ideas here, and refer to Supplementary Results B.2, B.1 for proofs of these claims.

For the first case when we drop the balancing condition, we can analyze the loci independently as there is no constraint on the entries of the ancestral profiles. Then, it suffices to observe that since the distance corresponds to the absolute difference, the function *c*(𝒯_*w*_; *x*) has a nice structure and we do not have to store an infinitely large dynamic programming table. When the integrality is removed, then, since both (i) and (ii) are *linear* constraints on the profiles and because *ℓ*_1_ norm minimization can be written as a linear program (LP), there is an LP formulation of the ZCNT small parsimony problem. As it is well known we can solve LPs in (weakly) polynomial time, this concludes the second case.

### 5.2 *Lazac* algorithm for ZCNT large parsimony

We develop a tree-search algorithm, *Lazac*, to find approximate solutions to the ZCNT large parsimony problem (Problem 2). Our procedure searches the space of copy number trees for a given copy number matrix *C* using sub-tree interchange operations [27] and relies heavily on the efficient algorithm we developed for the small parsimony problem (Problem 3) when the balancing condition (1) is dropped. The procedure is similar to the tree search procedure we developed for lineage tracing data [37]. Complete details on our tree search procedure are in Supplementary Methods A.1. *Lazac* is implemented in C++17 and is freely available at: github.com/raphael-group/lazac-copy-number.

## 6 Results

### 6.1 Comparison of copy number distances and phylogenies on prostate cancer data

We first investigated the differences between the CNT and ZCNT distances on copy number profiles inferred from bulk whole-genome sequencing data from ten metastatic prostate cancer patients [17]. We analyzed the copy number profiles for these patients published in [22]. For each pair of copy number profiles from distinct samples (e.g. anatomical sites) from the same patient, we computed the CNT distance *d*_CNT_ and ZNCT distance *d*_ZCNT_. We found that for all ten patients, the median relative difference |*d*_CNT_/*d*_ZCNT_ − 1| over all pairs of samples was less than 5% – and for most patients the relative difference was even smaller (Supplementary Figure 5, 6).

### 6.2 Evaluation on simulated data

We compared *Lazac* to several state-of-the-art methods for inferring copy number phylogenies – namely MEDICC2 [22], MEDALT [43], Sitka [35], and WCND [50] – on simulated data.

*Lazac* inferred the most accurate phylogenetic trees across varying number of cells (*n* = 100, 150, 200, 250, 600) and loci (*l* = 1000, 2000, 3000, 4000). In particular, we found that on all but one parameter setting, *Lazac* had the lowest median RF (Figure 2) and Quartet distance (Supplementary Methods A.5) on simulated instances (Supplementary Figure 1). Further, on large phylogenies containing *n* = 600 cells, *Lazac* showed an even larger improvement in median RF (Figure 2) and Quartet distance (Supplementary Methods A.5) over other methods, owing to its scalability. In terms of speed, *Lazac* was the fastest method on every instance, taking less than ∼ 250 seconds to run on the largest simulated dataset containing 600 cells. Further, it was ∼100 times faster than the other top performing methods Sitka and MEDICC2 (Figure 2b).

**Figure 2:**
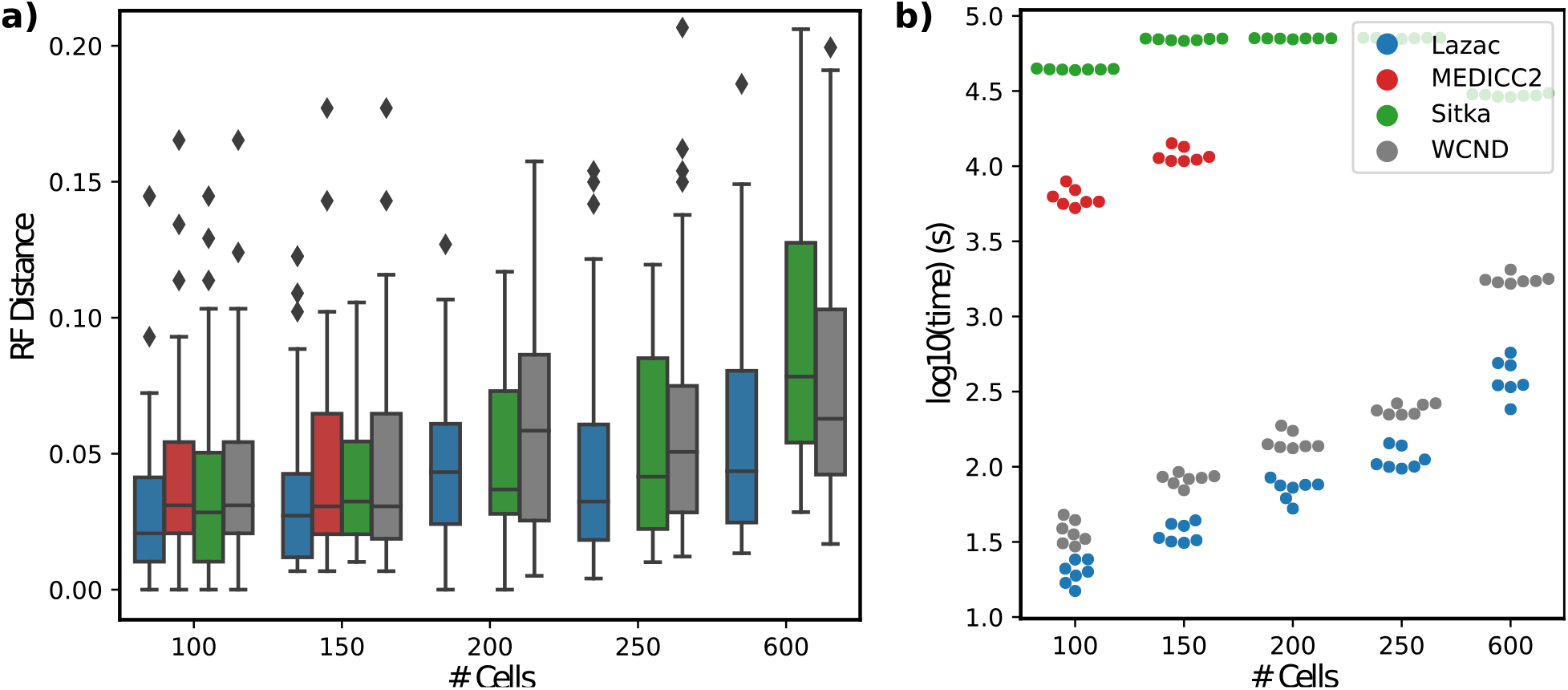
**(a)** Comparison of reconstruction accuracy (RF distance) on simulated data for several state-of-the art methods for copy number tree reconstruction with varying number of cells *n* = 100, 150, 200, 250, 600 across four sets of loci *l* = 1000, 2000, 3000, 4000 and seven random seeds *s* = 0, 1, 2, 3, 4, 5, 6. **(b)** Timing results for varying number of cells *n* = 100, 150, 250, 600 and fixed number of loci *l* = 4000. As MEDICC2 was too slow to run on more than 150 cells (with a 2 hour time limit), we exclude it from comparisons where the number of cells *n >* 150.

As a further evaluation of the differences between the ZCNT and CNT distances, we compared the trees obtained using distance-based phylogenetic methods with the ZCNT and CNT distances. Specifically, we compared the performance of applying neighbor joining on the ZCNT distances, referred to as *Lazac-NJ*, to three distance-based methods for reconstructing copy number phylognies: MEDICC2 [22], WCND [50], and MEDALT [43] on simulated data. MEDICC2 and WCND compute distances based on extensions of the CNT model and then apply neighbor joining to infer phylogenies. As such, they allow for a natural benchmark with which to compare our simpler, ZCNT distance. *Lazac-NJ* had nearly identical (within 1%) median RF and Quartet distance compared to other distance based methods (Supplementary Figure 2, 3) and was often the top performer. This provides evidence that even by itself, the ZCNT distance is useful for phylogenetic reconstruction.

### 6.3 Single-cell DNA sequencing data

We used *Lazac* to analyze single-cell whole genome sequencing (WGS) data from 25 human breast and ovarian tumor samples [15]. This dataset was generated using the DLP+ [23], single-cell whole-genome sequencing technology which produces ≈0.04× coverage from a median (resp. mean) of 636 (resp. 1457) cells per sample. The original study used Sitka [35], a method that uses the breakpoints between copy number segments as phylogenetic markers, to construct copy number phylogenies using this data.

We found that the phylogenies inferred by *Lazac* are substantially different than the phylogenies constructed by Sitka. Specifically, the normalized RF distance between pairs of phylogenies was greater than 0.90 in all cases (Supplementary Figure 9). In many cases, the normalized RF distance was 1, indicating that the phylogenies completely disagree.

To investigate these differences, we analyzed the concordance between the phylogenies and the assignments of cells to copy number clones, reported in the original publication [15]. Specifically, we defined the *clonal discordance score* as the parsimony score of the clonal labeling; i.e. the minimum number of changes in clone label that are required to label the leaves with the published clone labels. Thus, for a dataset with *k* clone labels the minimum possible clonal discordance score for a tree is *k* − 1 corresponding to the case each clone label is a clade in the tree. We find that on 18/25 of the samples, the *Lazac* phylogenies had substantially lower clonal discordance scores than the Sitka phylogenies (Supplementary Figure 4, 10) showing that the *Lazac* phylogenies were more concordant with the copy number clones compared to the published phylogenies. As a qualitative example, the phylogeny constructed for sample SA1184 by *Lazac* also appears more concordant with the copy number clones than that inferred by Sitka (Figure 3). Further details on the clonal discordance analysis are in Supplementary Methods A.3.

**Figure 3:**
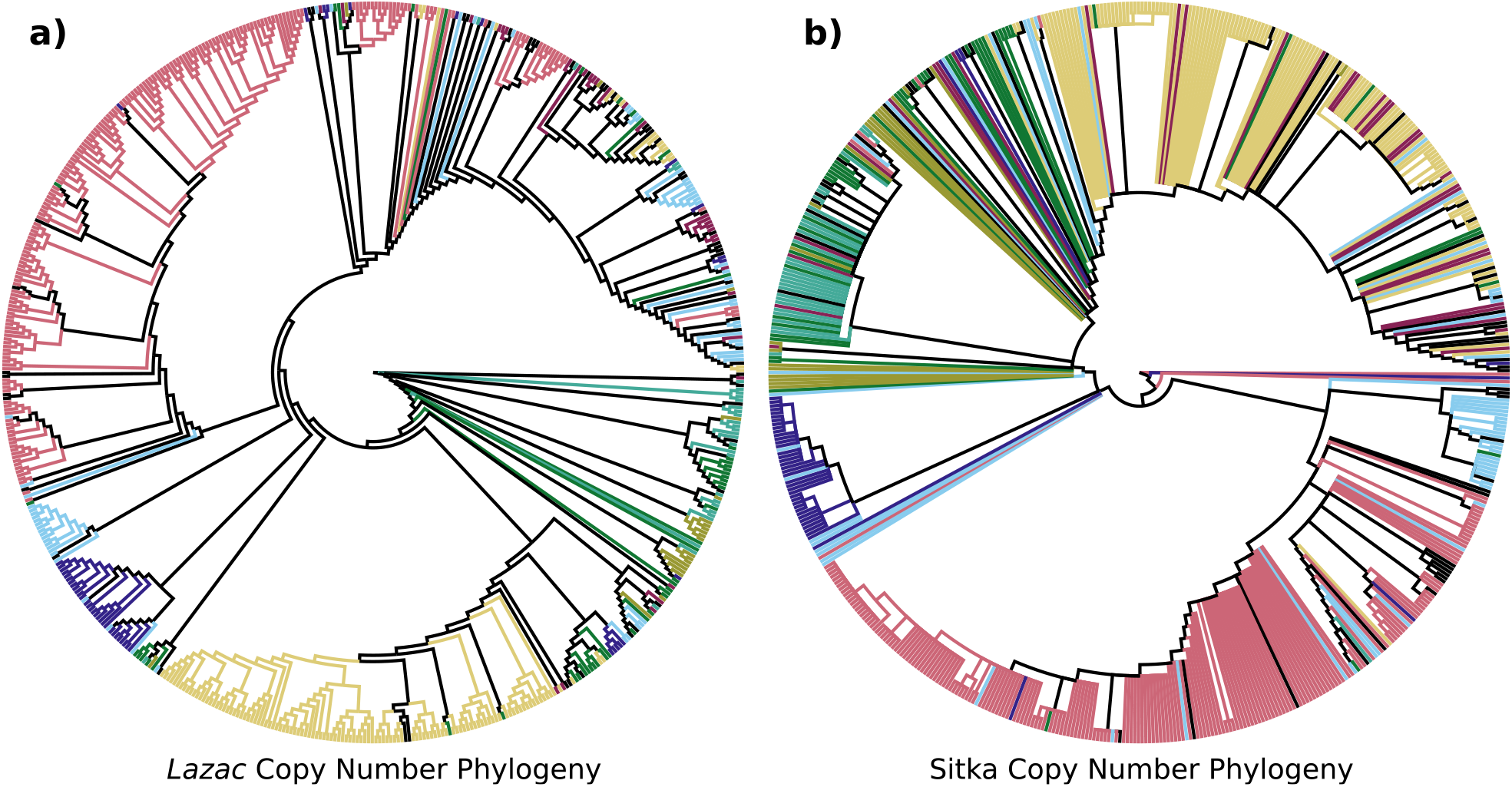
The copy number phylogenies inferred by *Lazac* **(a)** and Sitka **(b)** on sample SA1184 with the leaves colored by the corresponding clone labels, as visualized using Iroki [29]. The normalized RF distance between the two trees is 0.9869.

As a further evaluation of the *Lazac* and Sitka phylogenies, we examined whether somatic singlenucleotide variants (SNVs) supported the splits in each phylogeny, following the approach of [48]. Note that these SNVs were not used in the inference of either phylogeny, and thus they provide independent validation of the phylogeny. Given the extremely low sequence coverage (0.04× per cell), it is not possible to reliably measure SNVs of individual cells. Thus, we performed this analysis on the three samples (SA039, SA604, SA1035) with the largest number of cells. We identified subtrees in the phylogeny with at least 5% and at most 15% of the cells and identified SNVs present in the subpopulation of cells in these subtrees. Following the approach in [48], we perform a permutation test to determine whether the subtree is supported by more SNVs than expected (Supplementary Methods A.4). For all three samples, we found that the *Lazac* phylogenies had a greater fraction of supported subtrees (*P <* 0.05) than the *Sitka* phylogenies (Supplementary Figure 11). On the largest sample, SA1035, we identified five out of six supported subtrees (supported by 3175, 3334, 3799, 3435, and 3402 SNVs) for the *Lazac* phylogeny compared to only three of eight statistically significant subtrees (supported by 3426, 3129, and 3362 SNVs) for the Sitka phylogeny.

### 6.4 Approximation error of ZCNT small parsimony relaxations

We investigated the approximation error produced by the relaxations (Section 5.1) used in our two polynomial time algorithm’s for the ZCNT small parsimony problem. To perform this investigation, we first generated a set of 200 copy number phylogenies by stochastically perturbing a phylogeny inferred by Sitka [35] from single-cell whole genome sequencing data (Section 6.3). Then, for each phylogeny, we computed the optimal solution to the ZCNT small parsimony problem and its two relaxations using (integer) linear programming.

Importantly, we found that the exact solution to the ZCNT small parsimony problem and the solution obtained by relaxing the integrality condition were identical in every case. This leads us to believe that the relaxed linear program has a special structure, which we state as the following conjecture:

#### Conjecture 1

*The constraint matrix A of the linear program obtained by relaxing the integrality condition of the ZCNT small parsimony problem is totally unimodular*.

If true, this would imply that ZCNT small parsimony can be solved *exactly* in polynomial time using linear programming.

In contrast, dropping the balancing condition resulted in solutions with a lower score, implying that the balancing condition does meaningfully constrain the solution space. Specifically, the Pearson correlation between the score produced by dropping the balancing condition and the exact ZCNT small parsimony score was *R*^2^ = 0.972 (*p <* 10^−100^) across the 200 copy number phylogenies (Supplementary Figure 7).

Further, as the number of stochastic perturbations increased, both the parsimony score of the relaxed and the exact solutions increased (Supplementary Figure 8). This serves as a sanity check for the ZCNT small parsimony score since the phylogeny generally becomes further from ground truth as the number of stochastic perturbations increase.

## 7 Discussion

We introduced the *zero-agnostic copy number transformation* (ZCNT) model, a relaxation of the CNT model that allows for modification of zero copy number regions. We derived a closed-form expression for the ZCNT distance and presented polynomial time algorithms to solve two natural approximations of the small parsimony problem for copy number profiles. We used our efficient algorithm for the small parsimony problem to derive a method *Lazac*, to solve the large parsimony problem for copy number profiles. We demonstrated that on both real and simulated data, *Lazac* found better copy number phylogenies than existing methods.

There are multiple directions for future work. First, the complexity of the small parsimony problems for both the CNT and ZCNT models remains open, though we conjecture, and provide empirical evidence, that the latter is polynomial. Second, the algorithm we developed for the ZCNT large parsimony problem relies on a simple, hill climbing search using nearest-neighbor interchange operations. We expect that a more advanced approach that uses state-of-the-art techniques from phylogenetics [27, 32] could substantially improve both inference speed and accuracy. Finally, is to apply *Lazac* to other large single-cell WGS datasets [48, 28]. We anticipate that the scalability and accuracy of *Lazac* will be useful in analyzing the increasing amount of single-cell WGS cancer sequencing data.

## 8 Acknowledgements

We thank Uthsav Chitra and Gillian Chu for their helpful comments and review of this manuscript during several stages of its preparation. This work was supported by National Cancer Institute (NCI) grant U24CA248453 to B.J.R.

## A Supplementary Methods

### A.1 Large parsimony details

Our tree-search algorithm starts with an initial set of candidate trees *S* = (𝒯_1_, …, 𝒯_*k*_) and iteratively improves upon the trees by stochastic perturbations followed by a hill-climbing procedure. Specifically, at each iteration we select a candidate tree uniformly at random and perturb the tree using a random number *r* of nearest neighbor interchange (NNI) operations. With our perturbed candidate tree, we then perform local optimization using hill climbing to minimize the cost of the tree, where we use our small parsimony algorithm to efficiently evaluate the cost of each candidate topology. Once the hill-climbing procedure reaches a local minimum, we complete the iteration and update the candidate tree set if an improvement was found. The algorithm terminates after no improvement is found for a fixed number *I* of iterations.

For all experiments and analysis in this paper, the number of iterations prior to termination was set to *I* = 150. The number of random NNIs to perturb the candidate tree is selected uniformly at random from the discrete interval 0, {1, …, ⌊2.5*n*⌋} at every iteration. Our candidate tree set was generated by performing neighbor joining on the boundary insensitive distances and then randomly perturbing the the neighbor joining tree.

### A.2 Simulation details

We used a modified version of CONET’s [25] copy number phylogeny simulator. Specifically, we found that CONET’s simulation of tree structure was non-standard and opted to use a forward-birth death model [30] to simulate our topology. Once the tree structure was generated, we then used CONET’s simulator to sample copy number events on each vertex. We then took our event labeled copy number phylogeny and sampled the ground truth copy number states on the leaves of the phylogeny to obtain our copy number profiles.

To generate the tree topology, we used Cassiopeia’s [21] implementation of a forward-birth death model. We performed simulations for *n* = 100, 150, 200, 250, 600 leaves with a fitness parameter of 1.3 and an initial birth scale of 0.5. We drew the birth-death waiting times from an exponential distribution. With the topology, we randomly sampled events on each vertex using CONET with *l* = 1000, 2000, 3000, 4000 loci. We performed each simulation with parameters (*n, l*) a total of 7 times with unique random seeds *s* = 0, 1, 2, 3, 4, 5, 6. In total, there were 140 randomly simulated instances.

### A.3 Clonal concordance analysis

To analyze the concordance of the inferred phylogenetic trees with clonal information, we measured the minimum number of evolutionary events required to explain the clones. Specifically, for each sample clones were identified by clustering the GC-corrected read count profiles embedded using UMAP [26, 15]. The clone labels were then attached to the leaves of the inferred phylogenetic trees. With this clone labeled phylogenetic tree, we solved the small parsimony problem under the Wagner [42] model to obtain a parsimony score, *p*, which we call the *clonal discordance score*. This clonal discordance score is the minimum number of clonal transitions required to explain the cells of the phylogeny.

To compare across different phylogeny sizes, we computed the relative clonal discordance score between the *Lazac* and Sitka phylogenies as

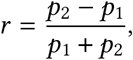

where *p*_1_, *p*_2_ are the clonal discordance scores of the *Lazac* and Sitka phylogenies respectively. In particular, a positive score indicates that the *Lazac* phylogeny is more concordant with the clones while a negative score indicates that the Sitka phylogeny is more concordant with the clones.

### A.4 Permutation test for analysis of SNV support

In this section we provide details about how subtrees of the phylogenies are identified for the analysis and the permutation test used for investigate if the subtrees are supported by the SNVs.

First, we describe how we identify subtrees in the phylogeny to analyze. Our goal is to identify subtrees that have enough cells so that pooling the reads from the cells and finding SNVs is feasible. However, if the number of cells is too large, the permutation test will not yield a significantly low p-value. To that end, we perform a breadth-first traversal of the nodes of the tree (starting from the root) to identify the desired subtrees. At each iteration, we compute the number of cells in the subtree rooted at the node (i.e. the number of leaves in the subtree). We select the subtree if (1) the number of cells in the subtree is more than 10% of the total number of cells (2) the number of cells in the subtree is less than 25% of the total number of cells and (3) the subtree is not contained in any of the subtrees selected in previous iterations.

Now, we provide details about the permutation test for a given subtree. We say that an SNV supports a subtree of a phylogeny if all the cells that yield a read harboring the SNV are contained in the subtree. We randomly permute the cell labels 500 times and count the number of SNVs supporting a given subtree. The p-value is empirically estimated by the ratio of the number of instances in which more SNVs support the subtree than with the original unpermuted cell labels with the total number of permutations tested (which is 500 in our study).

### A.5 Comparison to simulated trees

We assess the accuracy of the inferred trees compared to the ground truth simulated trees by employing two distinct tree dissimilarity metrics. These metrics are implemented in the TreeCmp tool [5] and the comparisons are done in a similar manner to the comparisons in our *Startle* [37] paper. Our metrics take a ground truth tree, 𝒯^*^, and an inferred tree, 𝒯, both in Newick format [6].

The Robinson-Foulds (RF) distance, *d*_RF_ (𝒯, 𝒯^*^), is a tree distance metric based on the induced bi-partitions in the input trees [33, 4]. Each edge *e* ∈ 𝒯 is associated with a bi-partition 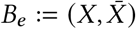 of its leaves, using the equivalence relation *x* ∼ *y* if *x* is connected to *y* in 𝒯_−*e*_, the forest formed by removing edge *e*. The set of bi-partitions for a tree 𝒯 is Bip(𝒯) = {*B*_*e*_ : *e* ∈ *E*(𝒯)}. The RF distance is then:

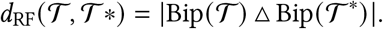

Similarly, the quartet distance, *d*_Q_(𝒯, 𝒯 ^*^), is a tree distance metric based on the induced quartets in the input trees [33, 4]. We define the set of quartets *Q*(𝒯) as the set of all consistent 4-leaf sub-trees with the unrooted topology of 𝒯. Then,

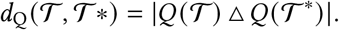

Finally, we used normalized versions of both *d*_RF_ and *d*_Q_ to enable comparison across different parameter settings. This normalization is implemented in TreeCmp [5] and described in their paper.

## B Supplementary Results

### B.1 ZCNT small parsimony: dropping the integrality condition

Let *Q* be the delta matrix obtained by applying the delta transformation to each row of the copy number matrix *M* (i.e. *q*_*ij*_ = Δ(*m*_*i*_)_*j*_). Using the formulation of the ZCNT small parsimony problem as stated in (Problem 3), we can write the objective as the following mathematical program.

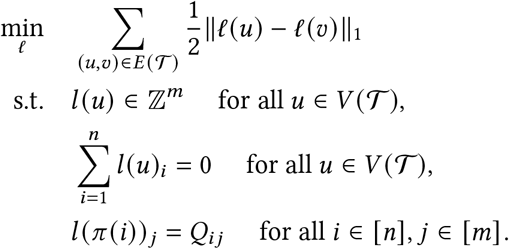

Notice that we can rewrite the optimization objective as a linear function subject to additional constraints. Specifically, 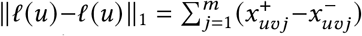 when 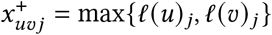 and 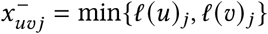.

And we can set 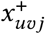 using the two linear constraints 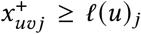 and 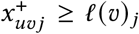; a similar procedure works for 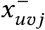. Then, by dropping the integrality condition *ℓ*(*u*) ∈ ℤ^*m*^, we obtain the following equivalent linear program.

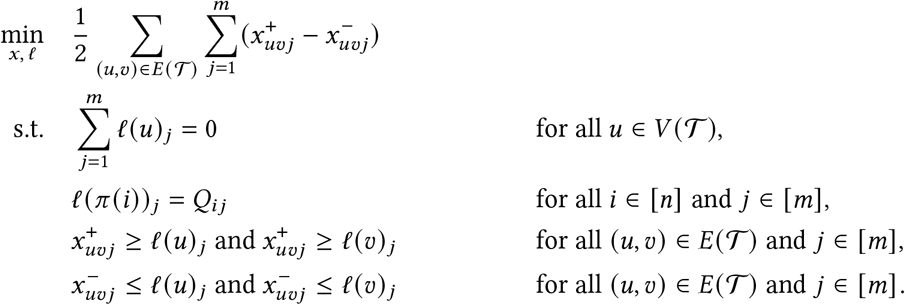

Since we can solve linear programs in (weakly) polynomial time, this proves the following theorem.

#### Theorem 4

*The ZCNT small parsimony problem can be solved in (weakly) polynomial time when the constraint that ℓ*(*u*) ∈ ℤ^*m*^ *is relaxed to ℓ*(*u*) ∈ ℝ^*m*^ *using a linear program with* 𝒪(*mn*) *variables and* 𝒪(*mn*) *constraints*.

### B.2 ZCNT small parsimony: dropping the balancing condition

When we drop the balancing condition, our problem becomes equivalent to the fixed topology rectilinear Steiner tree problem [36] on the delta profiles where the ancestral nodes lie in ℤ^*m*^. While there are several algorithms for the unrooted variant of this problem when Steiner vertices are in ℝ^*m*^ [13, 36, 9], our problem is different in that i) it assumes a rooted topology ii) the Steiner vertices are required to lie in ℤ^*m*^. Further, this problem has not been recently studied and we believe deserves a modern treatment. In this section, we present and prove the correctness of a linear time dynamic programming algorithm that solves the ZCNT small parsimony problem without the balancing condition.

We first observe that it suffices to analyze each locus independently. Let *Q* be the delta matrix obtained by applying the delta transformation to each row of the copy number matrix *M* (i.e. *q*_*ij*_ = Δ(*m*_*i*_)_*j*_). Let *ℓ*_*j*_ minimize the quantity ∑_(*u,v*) ∈*E*(𝒯)_ |*ℓ*_*j*_ (*u*) − *ℓ*_*j*_ (*v*) | and agree with the delta matrix *Q* on the leaves; that is, *ℓ*_*j*_ (*π*(*i*)) = *q*_*ij*_ for all cells *i*. Then, the labeling defined as *ℓ*(*u*) = (*ℓ*_1_(*u*), …, *ℓ*_*m*_ (*u*)) minimizes *J*(*ℓ*, 𝒯):

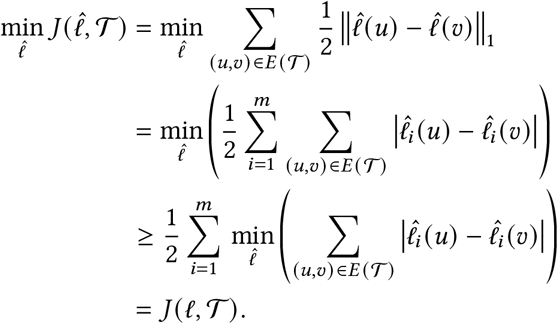

#### Algorithm 1 ZCNT small parsimony without the balancing condition

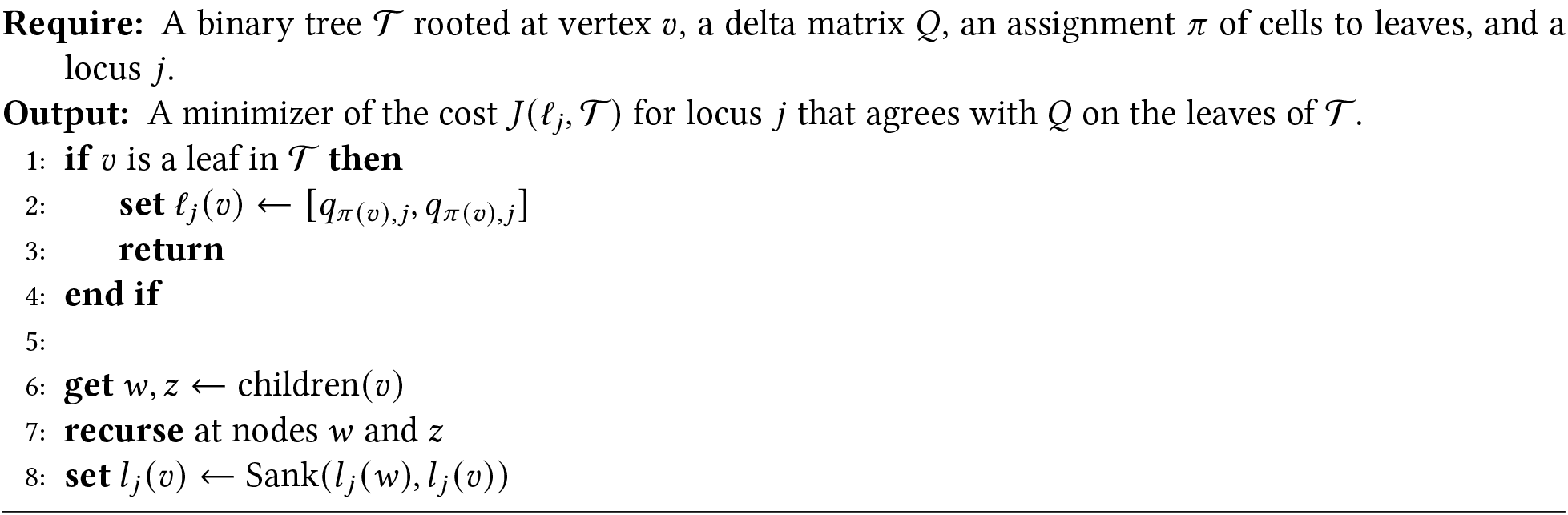

Thus, we can compute the cost of the optimal labeling *ℓ*_*j*_ for each locus independently and sum them together to obtain the entire cost. We also introduce the quantity *c*(𝒯 ; *x*) as the cost of the optimal labeling 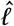 of 𝒯 that agrees with *ℓ* on the leaves of 𝒯 and has root label *x*. Our algorithm relies on the following easy to compute function on discrete intervals denoted [*a, b*] = {*a, a* + 1, …, *b* − 1, *b*}:

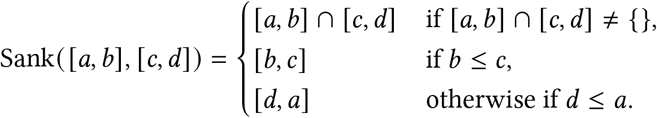

Our algorithm then applies the Sank (·, ·) function in a top-down recursive fashion, to compute *the interval* of optimal root labelings for each node. Though this procedure is quite natural, its proof of correctness is not immediately obvious and relies on a technical (Lemma 3).

#### Theorem 5

*(Algorithm 1) solves the ZCNT SCNP problem in* 𝒪(*n*) *time for a single locus when the balancing condition is dropped*.

*Proof*. Follows from induction on the size of the tree using (Lemma 3).

To prove the correctness of our procedure, we introduce the function dist(*x*, [*a, b*]) defined as the distance from *x* to the discrete interval [*a, b*]:

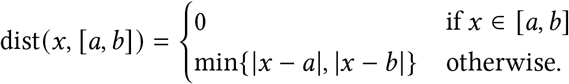

The correctness of our algorithm relies on two technical lemmas whose proofs are in (Section C).

#### Lemma 1

*If a* ≤ *b < c* ≤ *d are integers, then for all integers x the following inequality holds:*

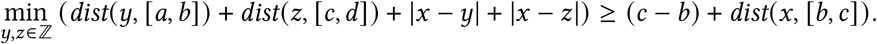

#### Lemma 2

*If a* ≤ *b, c* ≤ *d, a* ≤ *c, and b* ≥ *c are integers, then for all integers x the following inequality holds:*

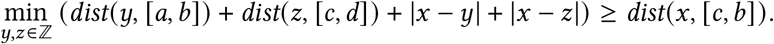

Then, the correctness of our algorithm follows from induction using the following lemma.

#### Lemma 3

*Let 𝒯 be a tree whose root vertex v has two children v*_1_ *and v*_2_. *Let* 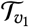 *and* 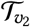 *denote the sub-trees rooted at v*_1_ *and v*_2_ *respectively. Then, suppose that*

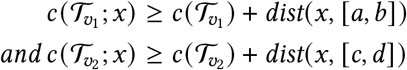

*where* [*a, b*] *and* [*c, d*] *are the set of optimal root labelings for* 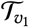 *and* 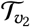. *Then*, [*e, f*] = *Sank*([*a, b*], [*c, d*]) *is the optimal set of root labelings for* 𝒯 *and*

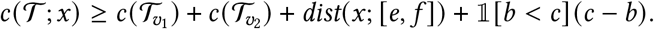

*Proof*. Without loss of generality we can assume that *a* ≤ *c*. Otherwise, we can swap the names of vertices *v*_1_ and *v*_2_. Let *x* be the labeling of the root of 𝒯. Then,

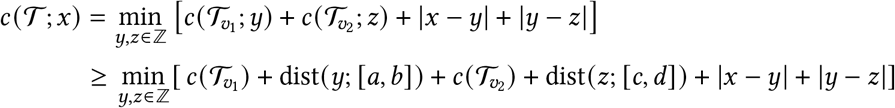

where the first equality follows from the definition of *c*(·; ·) and the fact that the distance is *ℓ*_1_. And the first inequality follows from our assumptions about 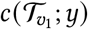 and 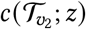.

We now consider the two cases of Sank([*a, b*], [*c, d*]) separately.

#### Case 1

*b < c*. In this case, we know from definition of Sank() that Sank ([*a, b*], [*c, d*]) = [*b, c*]. We want to show that

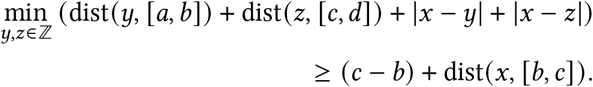

This will prove the desired inequality of the theorem. Then, to see that [*b, c*] is the optimal labeling of the root it is enough to observe that the inequality is realized when *x* ∈ [*b, c*]. As the proof of this inequality is rather technical and unenlightening, it is summarized in in (Lemma 1) and proven in Supplementary Proofs.

#### Case 2

Sank([*a, b*], [*c, d*]) = [*c, b*] and *c* ≤ *b*. In this case, we want to show that

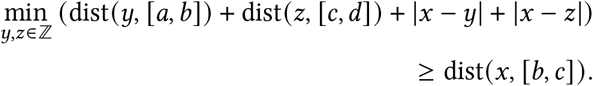

Which will again prove the desired inequality of the theorem. Then, to see that [*c, d*] is the optimal labeling of the root it is enough to observe that the inequality is realized when *x* ∈ [*c, d*]. The proof of this inequality is given in (Lemma 2).

## C Supplementary Proofs

### Theorem 1

*Let c*_*s,t,b*_ *be a zero-agnostic copy number event and δ*_*s,t,b*_ *be a delta event. Then*,

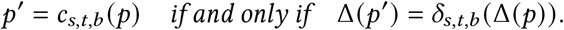

*Proof*. (⇒) Let *p*′ be the result of applying the zero-agnostic copy number event *c*_*s,t,b*_ to the profile *p*. Then, *i, i* − 1 ∈ {*s*, …, *t*}

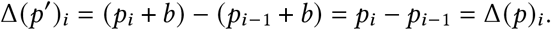

Similarly, if *i, i* − 1 ∉ {*s*, …, *t*}

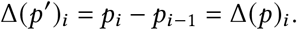

The remaining two cases occur when either *i* = *s* or *i* − 1 = *t*.

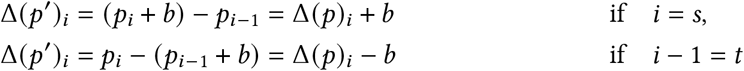

Thus, Δ(*p*′) is the result of applying the delta event *δ* to the profile Δ(*p*).

(⇐) This case is handled symmetrically.

### Proposition 3

*d*′(*u, v*) *is a distance metric. Further*,

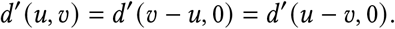

*Proof*. To see that *d*′(·, ·) satisfies the triangle inequality, it suffices to observe that the composition of delta sequences taking *u* to *v* and *v* to *w* transforms *u* to *w*. It is clearly reflexive since no delta event needs to be applied to map *u* to itself.

To see symmetry and the above equality, we observe that something stronger holds. Let

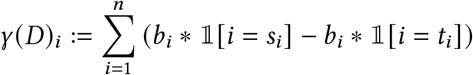

be the net change of coordinate *i* induced by the delta sequence *D* = (*δ*_1_, …, *δ*_*n*_) where *δ*_*i*_ = (*s*_*i*_, *t*_*i*_, *b*_*i*_). Then, *D* takes *u* to *v* if and only if *γ*(*D*)_*i*_ = *u*_*i*_ − *v*_*i*_ for all *i* ∈ {1, …, *n*}. This follows from the definition of our delta event and the fact that applying *D* to *u* results in a profile *v* defined by its entries as *v*_*i*_ = *u*_*i*_ + *γ*(*C*)_*i*_.

### Proposition 4

*For a vector p*′ = Δ(*p*), *the sum of the magnitudes of the positive entries equals the sum of the magnitudes of the negative entries. That is*,

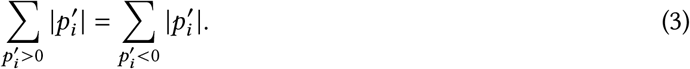

*Proof*. We first notice that 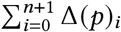 expands to a telescoping sum after applying our definition of a delta profile. Specifically,

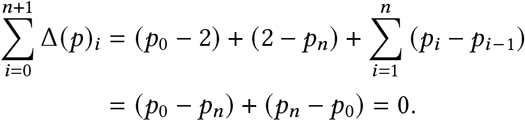

Then, we rewrite the left hand side of the sum

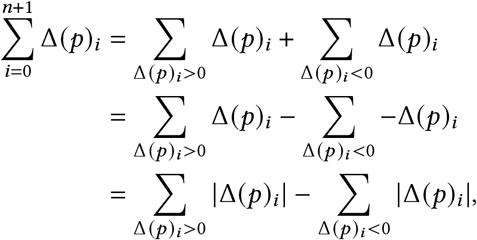

where the last equality follows from the definition of absolute value.

### Proposition 1

*The delta map* Δ : ℤ^*n*^ → 𝒟_*n*+1_ *is invertible*.

*Proof*. By (Proposition 4) the range of the map Δ lies in the space of vectors in ℤ^*n*+1^ satisfying the balance condition. Thus, it suffices to show that Δ is injective and can reach all vectors (i.e. is surjective) in this space.

To see that the map is injective, let *x, y* ∈ ℤ^*n*^ be two distinct vectors. Then, define *i* as the first coordinate in which these vectors differ. Since *i* is the first coordinate in which these vectors differ, *x*_*i*−1_ = *y*_*i*−1_ but *x*_*i*_ ≠ *y*_*i*_. Thus,

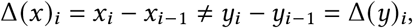

proving that Δ is injective.

To see the map is surjective, let *y* ∈ ℤ^*n*+1^ be a vector satisfying the balance condition. Define a vector *x* ∈ ℤ^*n*^ such that *x*_0_ = *y*_0_ + 2, and *x*_*i*_ = *y*_*i*_ + *x*_*i*−1_ for *i* ∈ {1, …, *n*}. Then, Δ(*x*) agrees with *y* on the first *n* coordinates, but since *y* and Δ(*x*) both satisfy the balance condition by (Proposition 4), their last coordinate is also determined, proving that Δ(*x*) = *y*.

### Corollary 3

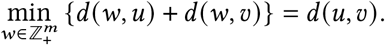

*Proof*. By the triangle inequality, *d*(*u, w*) + *d*(*w, v*) ≥ *d*(*u, v*) for all profiles *w*. Setting *w* = *u* or *w* = *v* achieves equality, proving that setting *w* to either *u* or *v* minimizes the expression *d*(*w, u*) + *d*(*w, v*) as *d*(*u, w*) = *d*(*w, u*).

### Lemma 1

*If a* ≤ *b < c* ≤ *d are integers, then for all integers x the following inequality holds:*

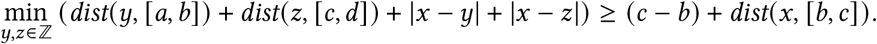

*Proof*. The statement follows from a case analysis on the locations of the variables we are optimizing over: *y, z*. However, before any case analysis, we prove that

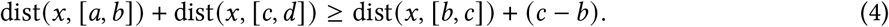

To see this, we perform a case analysis on *x*. If *x* ≤ *b* we have

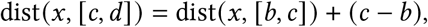

since *x* is to the left of the interval [*b, c*] which is to the left of the interval [*c, d*]. If *x* ≥ *c* we have

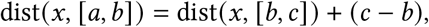

since *x* is to the right of the interval [*b, c*] which is to the right of the interval [*a, b*]. And finally, if *x* ∈ [*b, c*] then,

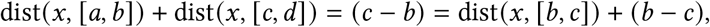

since dist(*x*, [*b, c*]) = 0.

### Case 1

*y* ∈ [*a, b*] and *z* ∈ [*c, d*]

In this case, dist(*y*, [*a, b*]) + dist(*z*, [*c, d*]) = 0, so

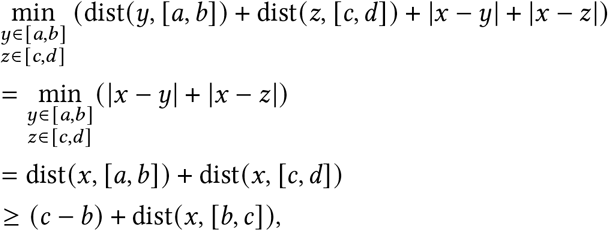

where the second equality follows from the fact that dist(*x*, [*a, b*]) = min_*y*∈[*a,b*]_ |*x* − *y*| and the inequality follows from (4).

### Case 2

*y* ∈ [*a, b*] and *z* ∉ [*c, d*] OR *y* ∉ [*a, b*] and *z* ∈ [*c, d*]

Since the two cases are symmetric, we only need to consider the former situation where *y* ∈ [*a, b*] and *z* ∉ [*c, d*]. In this case,

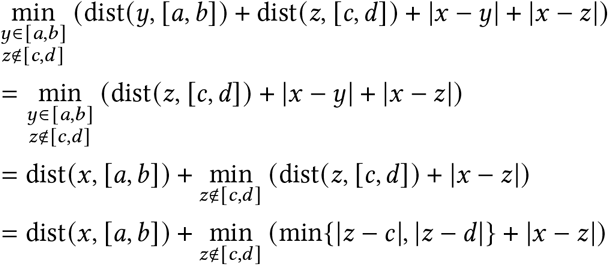

where the second equality follows the fact that dist(*x*, [*a, b*]) = min_*y*∈[*a,b*]_ |*x* − *y* | and the third equality from the definition of dist(·, ·). We now analyze two sub-cases separately.

**Sub-case 1:** *x* ∈ [*c, d*] In this case, dist(*x*, [*a, b*]) = dist(*x*, [*b, c*]) + (*c* − *b*) and we are done.

**Sub-case 2:** *x* ∉ [*c, d*]

Then,

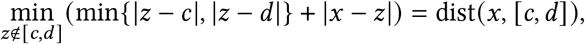

since the minimizer is found by setting *z* = *x*. From the above equality,

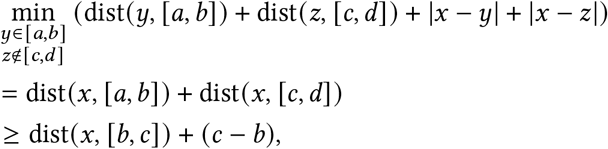

where the inequality again follows from (4).

### Case 3

*y* ∉ [*a, b*] and *z* ∉ [*c, d*]

In this case, by the definition of dist(·, ·)

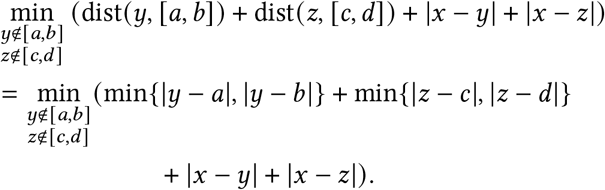

We now analyze two sub-cases separately. **Sub-case 1:** *x* ∉ [*a, b*] and *x* ∉ [*c, d*] Now, by the same reasoning as before,

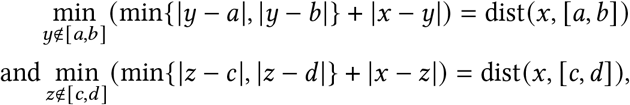

since the minimizer is found by setting *y* = *x* and *z* = *x*. Thus, by independence of the terms *y* and *z*,

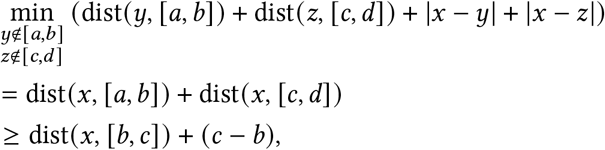

where the second inequality follows from (4).

**Sub-case 2:** *x* ∈ [*a, b*] OR *x* ∈ [*c, d*]

By symmetry, without loss of generality we assume that *x* ∈ [*a, b*]. Since the two intervals are disjoint *x* ∈ [*a, b*] implies *x* ∉ [*c, d*]. Then since *x* ∉ [*c, d*],

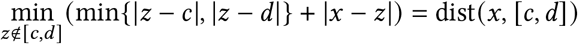

since the minimizer is realized by setting *z* = *x*. Finally since dist(*x*, [*a, b*]) = 0,

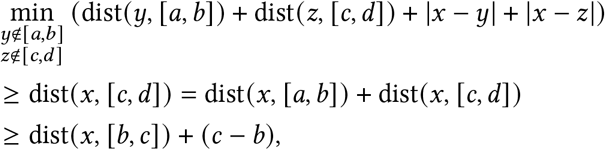

where the second inequality follows from (4). As this is the final case, we have proven the original claim.

### Lemma 2

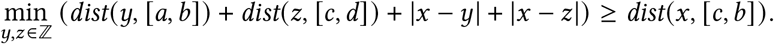

*Proof*. The proof again follows by a case analysis, but it is much simpler than in (Lemma 1).

### Case 1

*x* ∈ [*c, b*]

This case is trivial since dist(*x*, [*c, b*]) = 0 and the left hand side of the inequality is always non-negative.

### Case 2

*x* ≤ *c* or *x* ≥ *b*

As these cases are symmetric, it suffices to only consider the former case where *x* ≤ *c*. First, if *z* ≤ *c*,

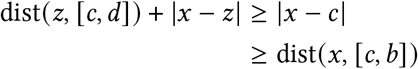

since the minimizer is found when *z* ∈ [*x, c*]. Second, if *z* ≥ *c*,

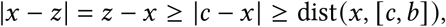

since *z* is to the right of *c* while *x* is to the left of *c*. This completes the proof.

## D Supplementary Figures

We have the following supplementary figures and tables:

- (Supplementary Figure 1) compares the reconstruction accuracy in terms of Quartet distance on CONET simulated data across state-of-the-art methods.
- (Supplementary Figure 2) compares the baseline reconstruction accuracy for a variety of extremely simple distance based methods including our distance method, *Lazac Nℱ*.
- (Supplementary Figure 3) compares the reconstruction accuracy on CONET simulated data for a variety of distance based methods including our distance method, *Lazac Nℱ*.
- (Supplementary Figure 4) compares the clonal discordance scores between *Lazac* and Sitka inferred phylogenies from an in vitro cell line system modeling instability in human and breast ovarian tumours.
- (Supplementary Figure 5) displays the relative difference between the ZCNT and CNT distance for patient 008 from a metastatic prostate cancer tumor sample [17].
- (Supplementary Figure 6) displays the relative difference between the ZCNT and CNT distance for patient 012 from a metastatic prostate cancer tumor sample [17].
- (Supplementary Figure 7) compares the solutions to the relaxed and unrelaxed ZCNT small parsimony problem across 200 phylogenies.
- (Supplementary Figure 8) compares the solutions to the relaxed and unrelaxed ZCNT small parsimony problem across 200 phylogenies as a function of the number of stochastic perturbations used to obtain that phylogeny.
- (Supplementary Figure 9) shows the distribution of normalized RF distance between trees inferred by *Lazac* and trees inferred by Sitka on human breast and ovarian tumour data [15].
- (Supplementary Figure 10) a table displaying the results of our concordance analysis on trees inferred by *Lazac* and trees inferred by Sitka on human breast and ovarian tumour data [15].
- (Supplementary Figure 11) Results of somatic SNV analysis on subset of samples from an in vitro cell line system modeling instability in human breast and ovarian tumours [15].

**Supplementary Figure 1:**
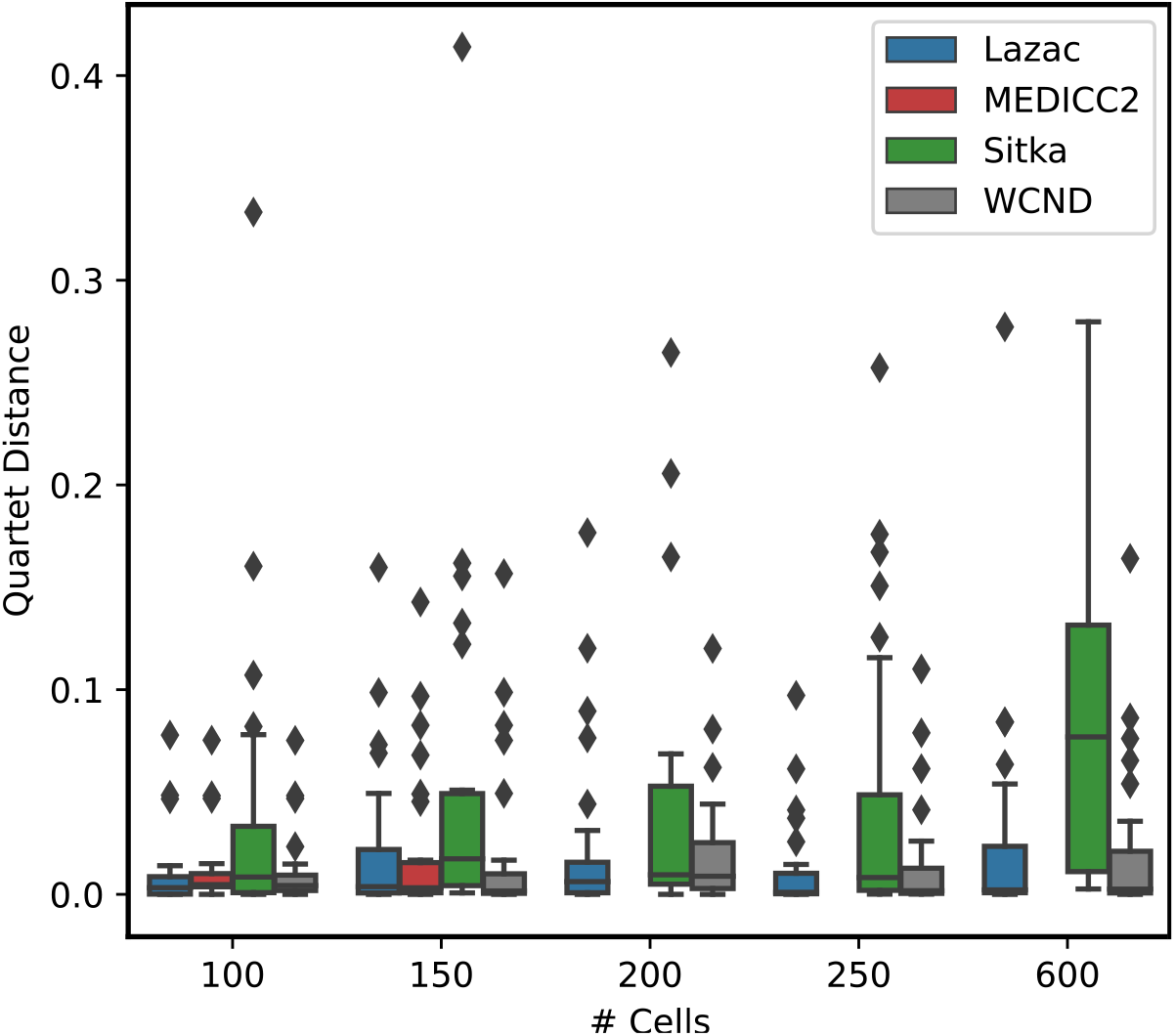
Baseline reconstruction accuracy (Quartet distance) on CONET simulated data for copy number tree reconstruction with varying number of cells *n* = 100, 150, 200, 250, 600 across four sets of loci *l* = 1000, 2000, 3000, 4000 and seven random seeds *s* = 0, 1, 2, 3, 4, 5, 6.

**Supplementary Figure 2:**
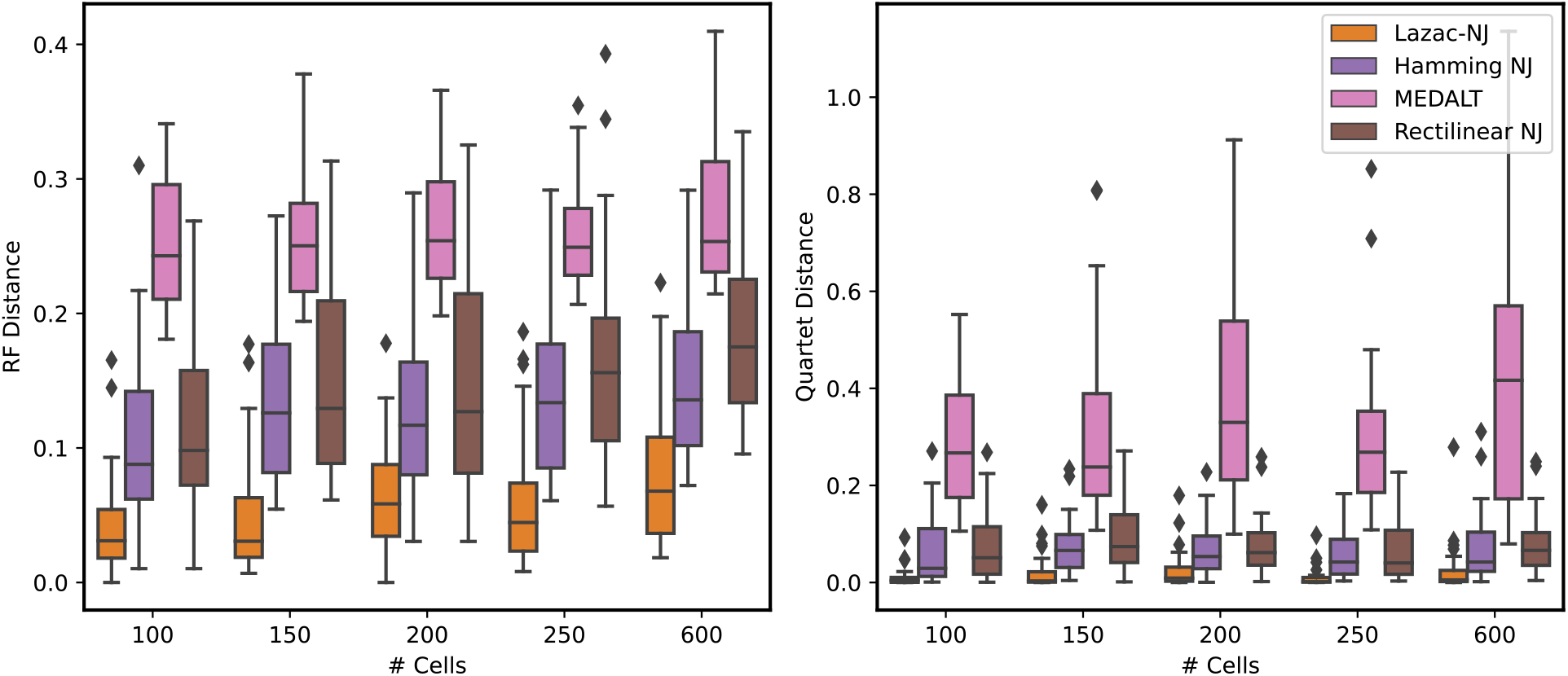
Baseline reconstruction accuracy on (left: RF distance; right: Quartet distance) CONET simulated data across simple NJ methods for copy number tree reconstruction with varying number of cells *n* = 100, 150, 200, 250, 600 across four sets of loci *l* = 1000, 2000, 3000, 4000 and seven random seeds *s* = 0, 1, 2, 3, 4, 5, 6.

**Supplementary Figure 3:**
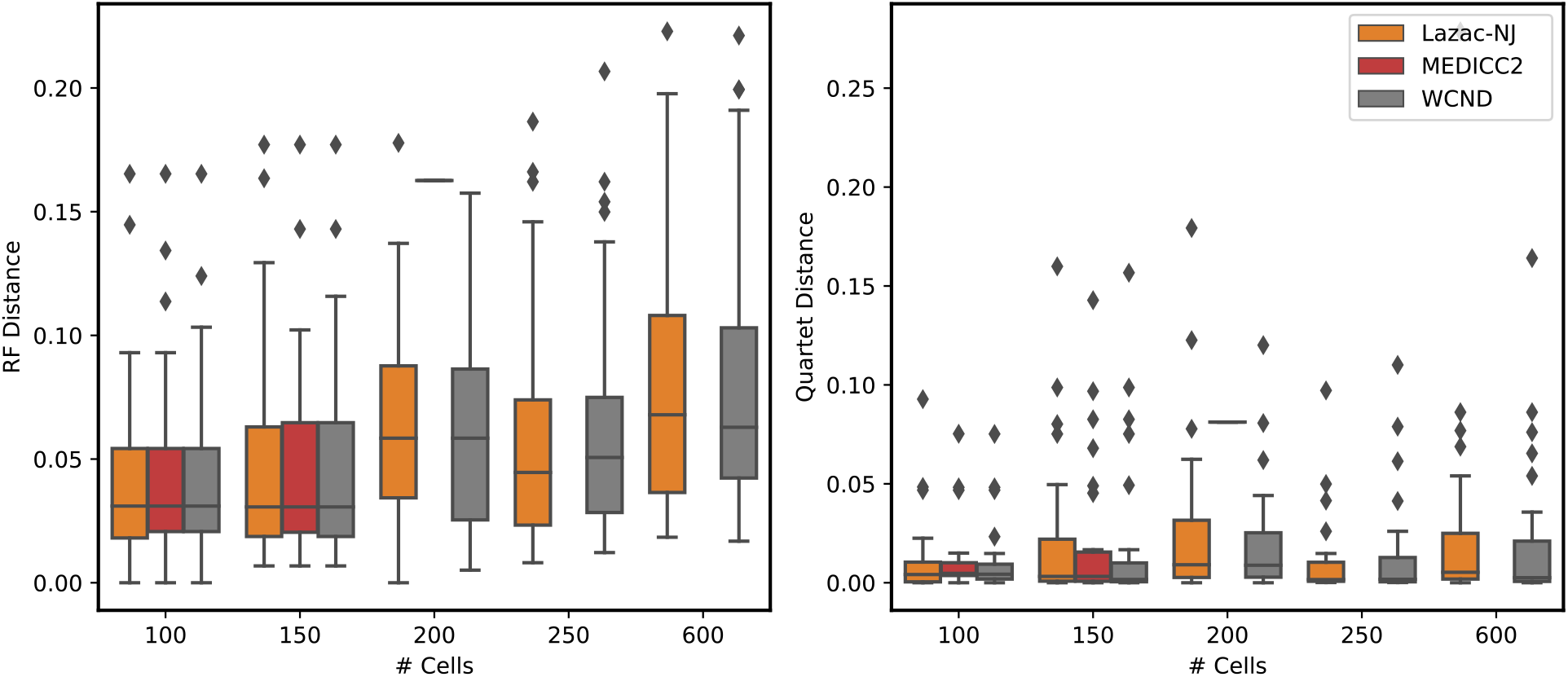
Comparison of reconstruction accuracy (left: RF distance; right: Quartet distance) CONET simulated data across *distance based* methods for copy number tree reconstruction with varying number of cells *n* = 100, 150, 200, 250, 600 across four sets of loci *l* = 1000, 2000, 3000, 4000 and seven random seeds *s* = 0, 1, 2, 3, 4, 5, 6. As MEDICC2 was too slow to run on more than 150 cells, we exclude it from comparisons where the number of cells *n >* 150.

**Supplementary Figure 4:**
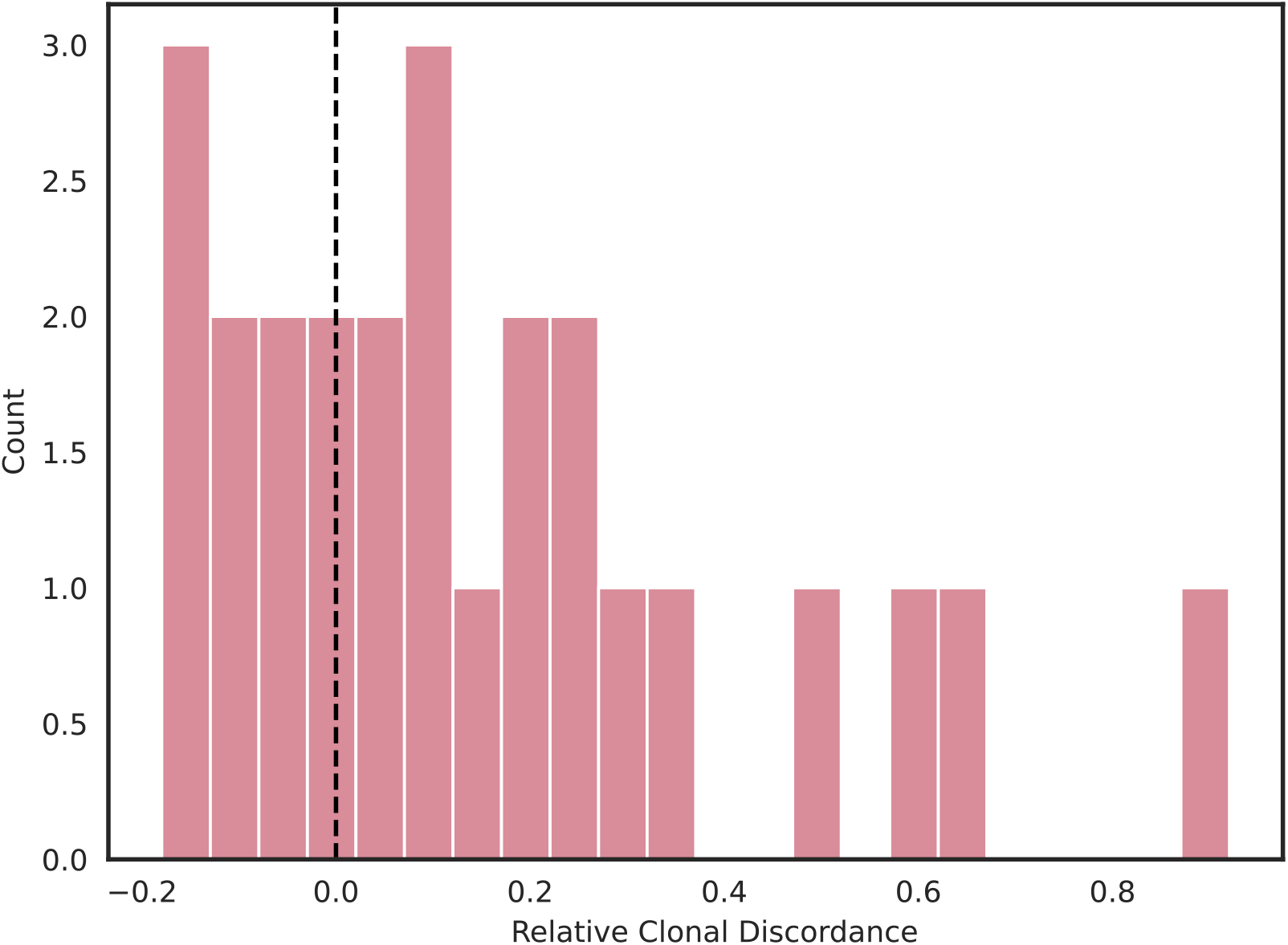
The relative clonal discordance score 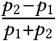 where *p*_1_, *p*_2_ are the clonal discordance scores of the *Lazac* and *Sitka* inferred phylogenies respectively.

**Supplementary Figure 5:**
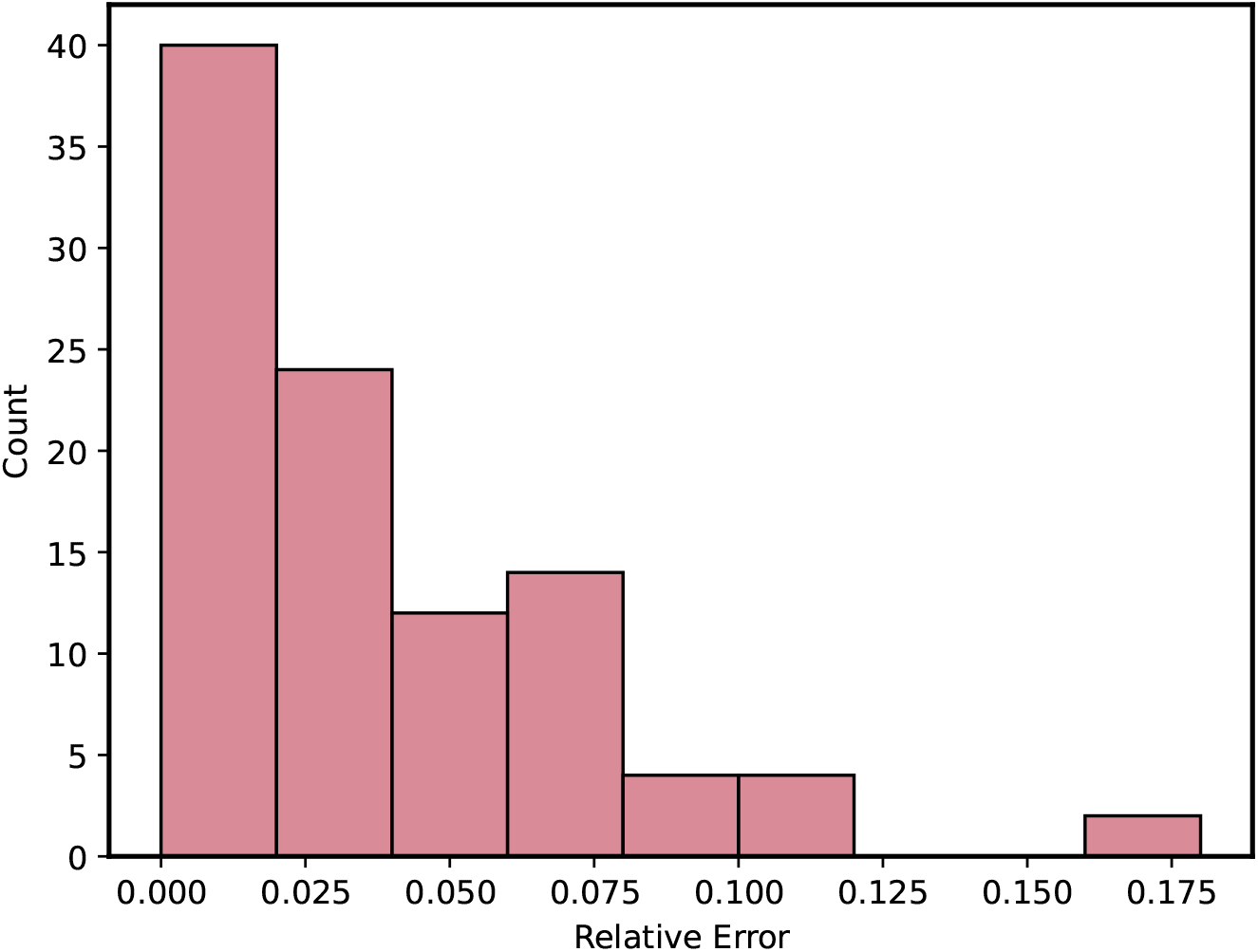
Relative difference between the ZCNT distance *d*(*p, p*′) and the CNT distance computed for patient 8 from a metastatic prostate cancer tumor sample [17].

**Supplementary Figure 6:**
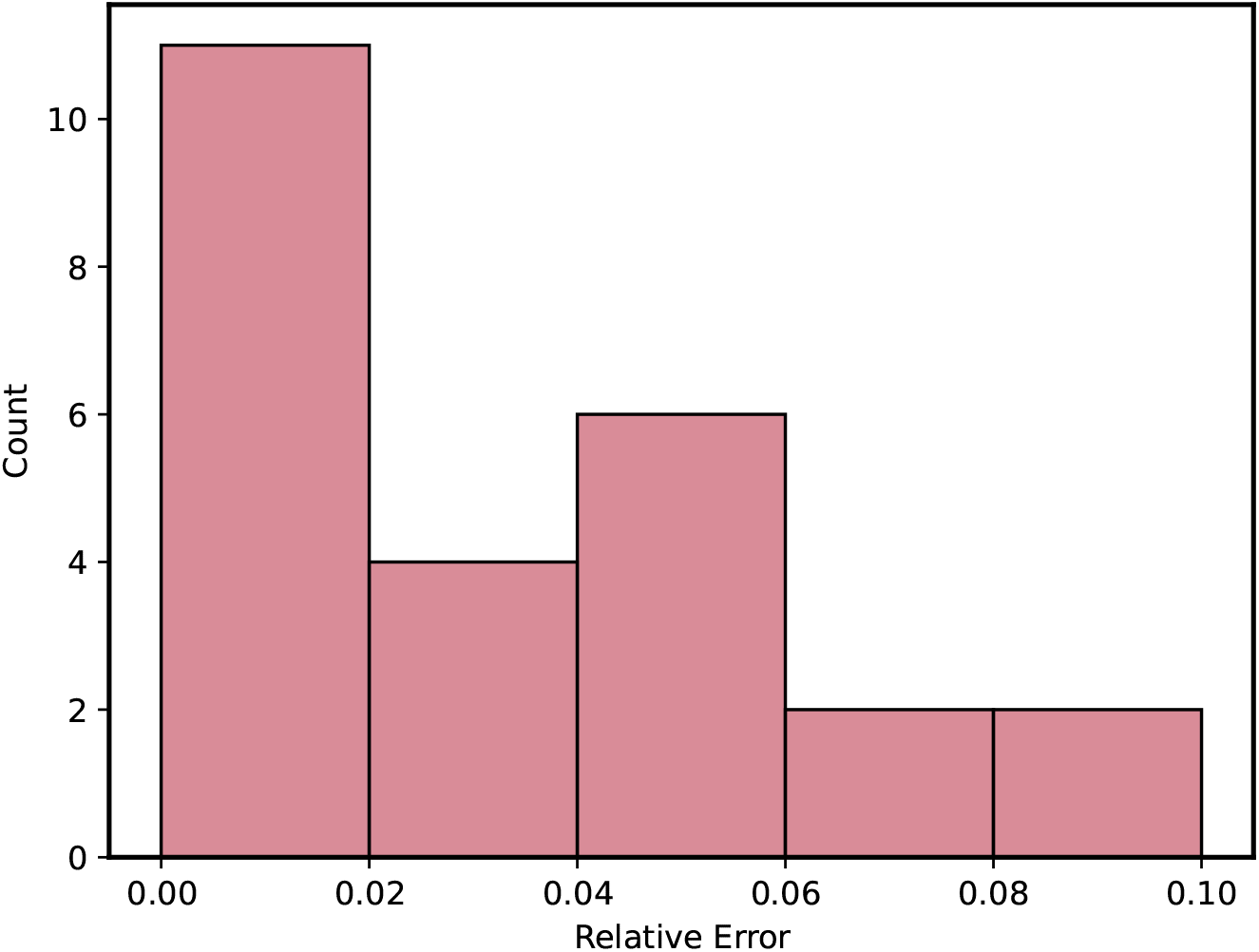
Relative difference between the ZCNT distance *d*(*p, p*′) and the CNT distance for patient 12 from a metastatic prostate cancer tumor sample [17].

**Supplementary Figure 7:**
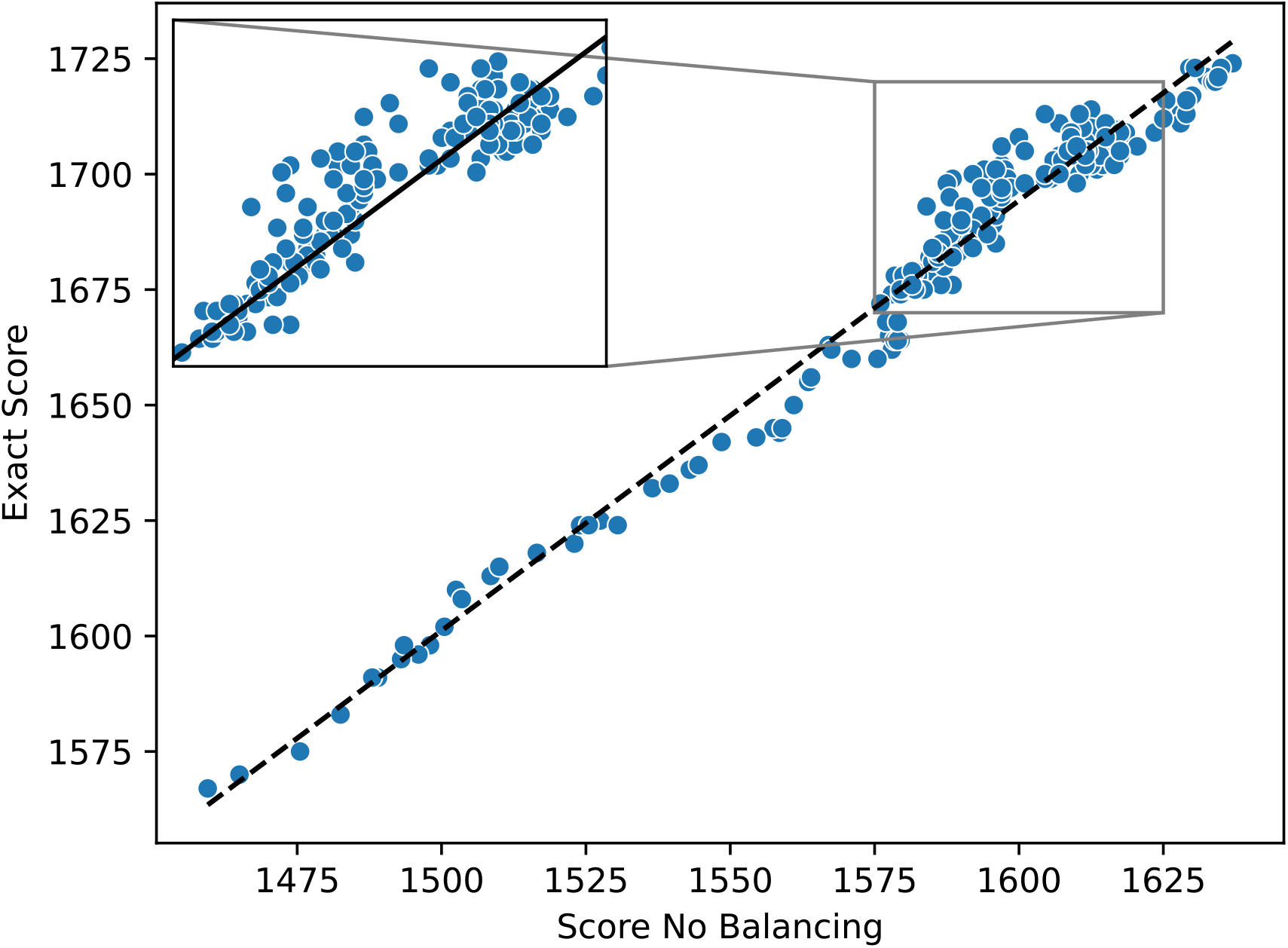
The exact versus the relaxed score of the optimal solution to the ZCNT small parsimony problem when the balancing condition is removed across 200 phylogenies. The 200 phylogenies were obtained by stochastic perturbation of the phylogeny inferred Sitka [35] on sample SA1053. The dotted line is computed by performing linear regression and is defined by *y* = 0.9313 * *x* + 204.1 with an *R*^2^ = 0.972 and a *p* = 1.05 * 10^−156^.

**Supplementary Figure 8:**
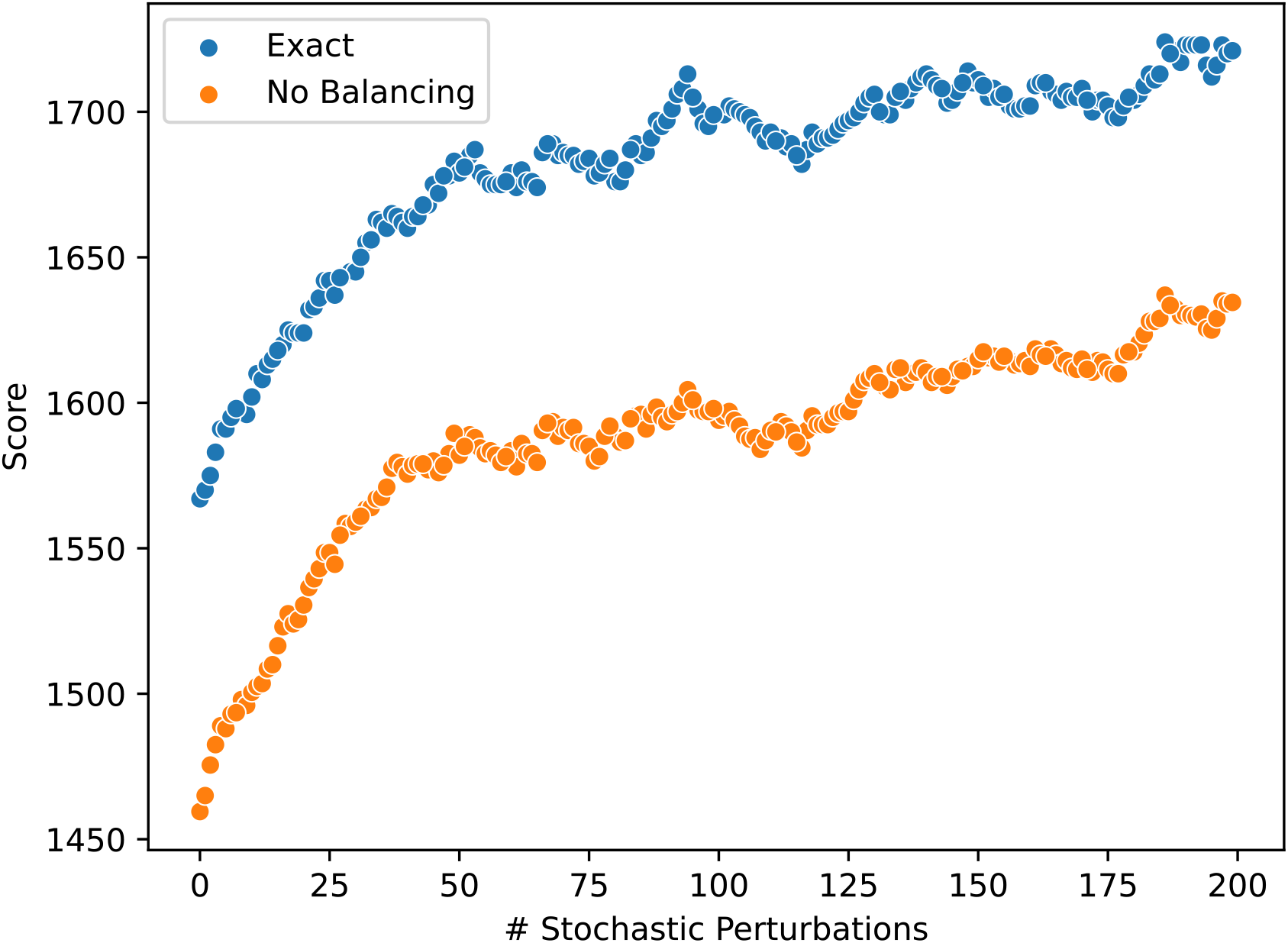
The exact and relaxed scores of the optimal solution to the ZCNT small parsimony problem as a function of the number of stochastic perturbations applied to the phylogeny inferred by Sitka [35] on sample SA1053.

**Supplementary Figure 9:**
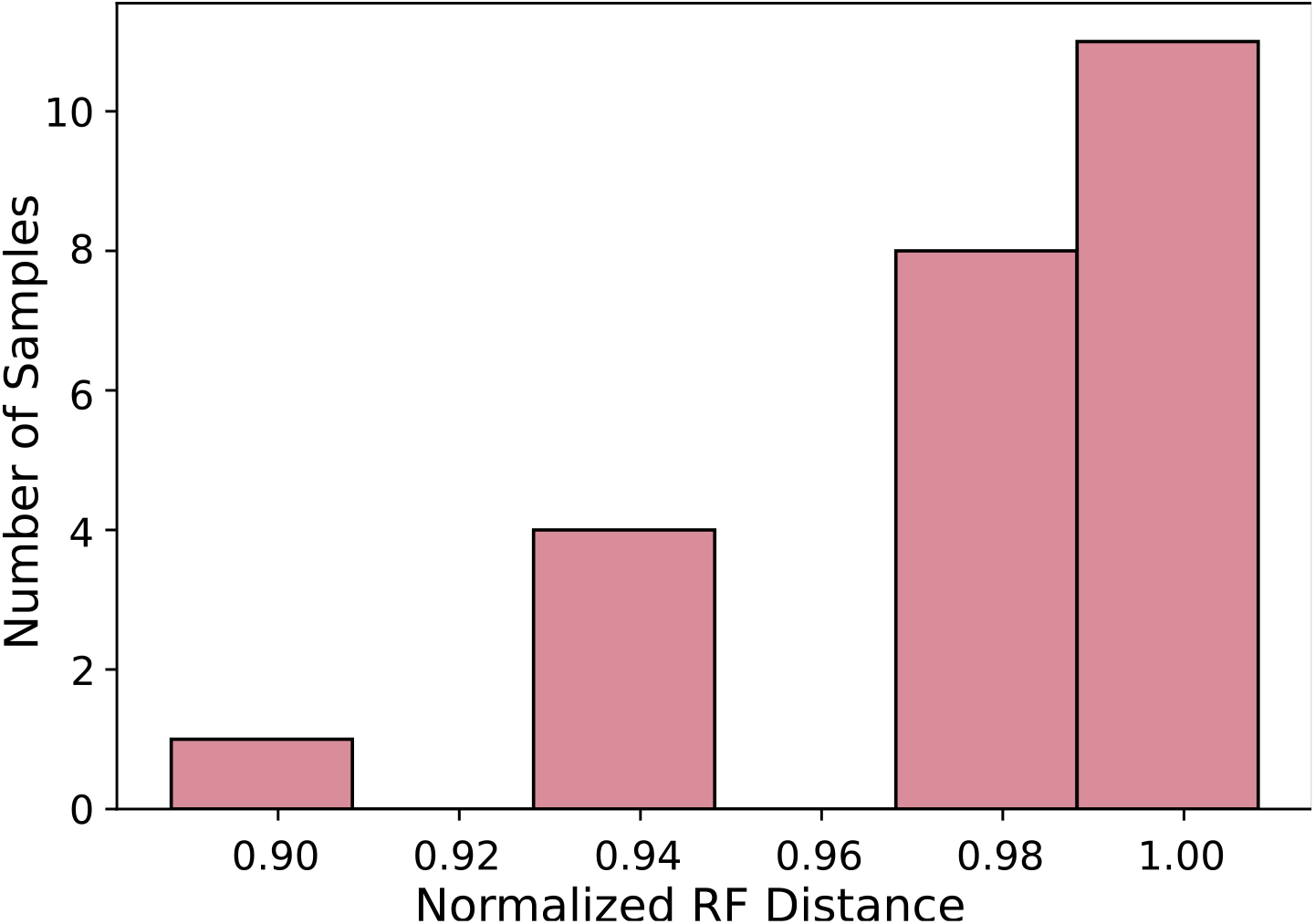
RF distances between *Lazac* and Sitka inferred trees on copy number profiles from human breast and ovarian tumour data [15].

**Supplementary Figure 10:**
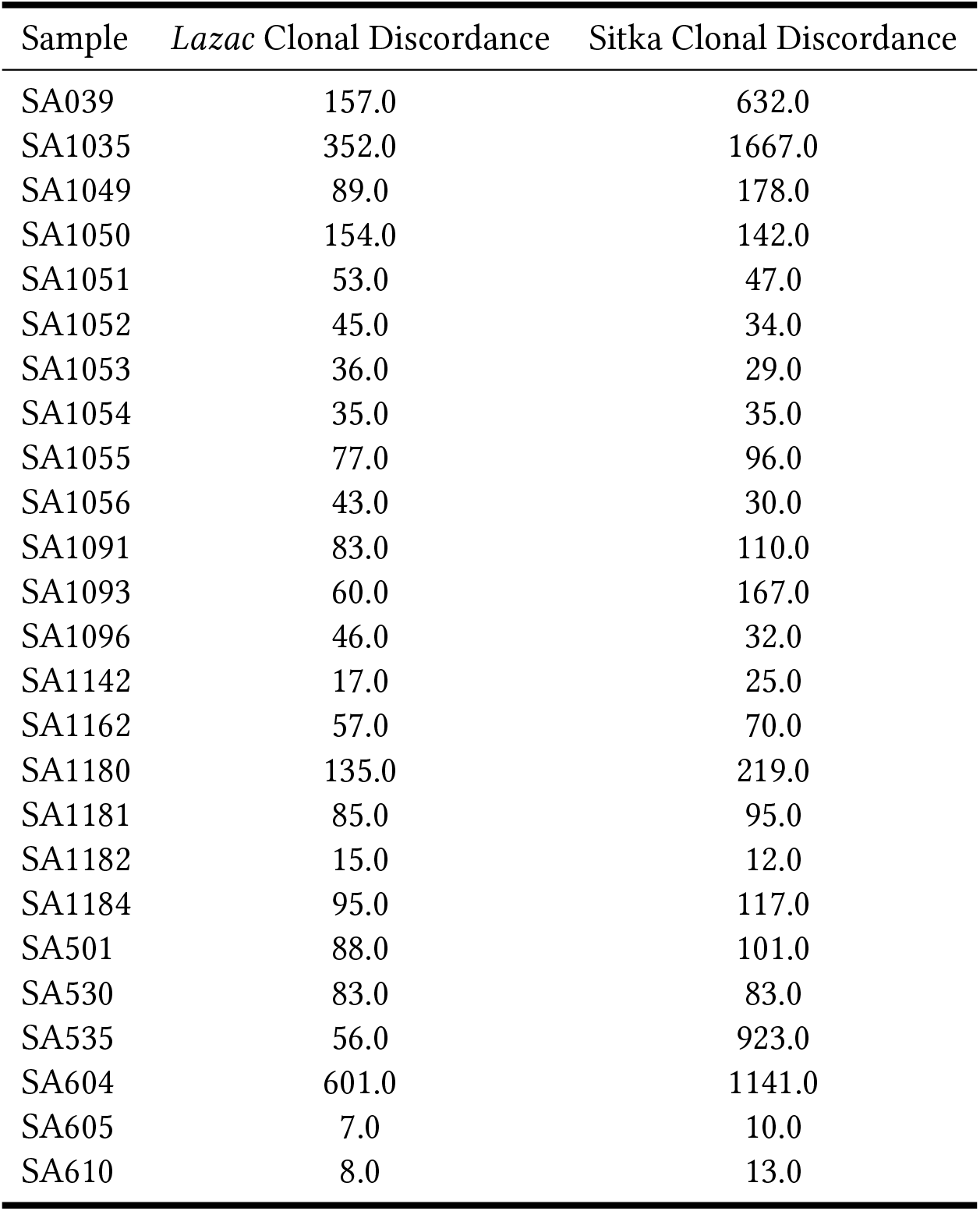
Comparison of *Lazac* and Sitka clonal discordance scores across 25 samples from an in vitro cell line system modeling instability in human breast and ovarian tumours [15].

**Supplementary Figure 11:**
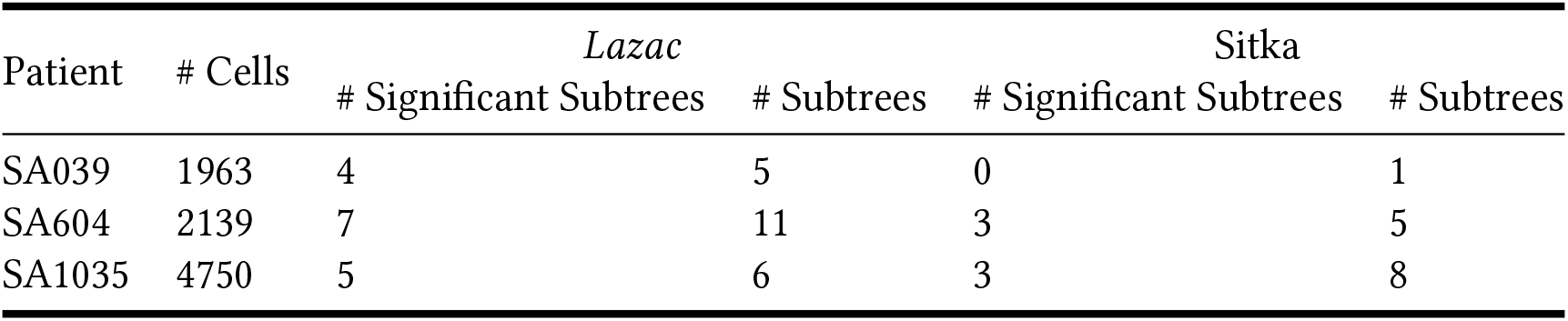
Results of somatic SNV analysis on subset of samples from an in vitro cell line system modeling instability in human breast and ovarian tumours [15]. Subtrees were called significant if the SNV permutation test p-value was below 0.05 for that clone.

